# Ribonucleotide reductase, a novel target for gonorrhea

**DOI:** 10.1101/2020.11.19.389957

**Authors:** J. Narasimhan, S. Letinski, S. Jung, A. Gerasyuto, J. Wang, M. Arnold, G. Chen, J. Hedrick, M. Dumble, K. Ravichandran, T. S. Levitz, C. Chang, C. L. Drennan, J. Stubbe, G. Karp, A. Branstrom

## Abstract

Antibiotic resistant *Neisseria gonorrhoeae (Ng)* are an emerging public health threat due to increasing numbers of multidrug resistant (MDR) organisms. We identified two novel orally active inhibitors, PTC-847 and PTC-672, that exhibit a narrow spectrum of activity against *Ng* including MDR isolates. By selecting organisms resistant to the novel inhibitors and sequencing their genomes, we identified a new therapeutic target, the class Ia ribonucleotide reductase (RNR). Activity studies and negative stain electron microscopy of the *Ng* Ia RNR suggest that these inhibitors potentiate conversion of its active α_2_β_2_ state to an inactive α_4_β_4_ similar to states first identified with the *Escherichia coli (Ec)* Ia RNR. Resistance mutations in *Ng* map to the *N*-terminal, ATP cone domain of its α subunit and disrupt the interaction with the β subunit required to form the specific quaternary inhibited state. Oral administration of PTC-672 reduces *Ng* infection in a mouse model and may have therapeutic potential for treatment of *Ng* that is resistant to current drugs.

## INTRODUCTION

Increasing resistance to current therapeutics against many pathogenic bacteria (Walsh 2015, Gerasyuto 2018) require new treatment options with unique mechanism(s) of action. Recently, we reported the discovery of a series of fused indolyl-containing 4-hydroxy-2-pyridones that showed inhibited bacterial DNA synthesis by targeting mutant DNA topoisomerases (gyrase, topoisomerase IV) and improved in vitro antibacterial activity against a range of fluoroquinolone resistant Gram negative strains (Gerasyuto 2018).

The interesting activities of a subset of 4-hydoxy-2-pyridones provided the impetus for synthesis of additional chemotypes with this core (Figure 1) and their effectiveness against additional pathogenic strains including *Ng* and *N. meningitidis (Nm)*. One World Health Organization (WHO) global challenge in 2012 (WHO 2012) focused on *Ng* isolates. They reported seventy eight million *Ng* infections worldwide with 90% of infections in low and middle income countries. In the US, during 2017–2018, the rate of reported *Ng* infections increased 5.0% (583,405 US reported cases in 2018 from CDC 2019a), and the rate increased among both sexes, in all regions of the US, and among all ethnicity groups. *Ng* was reported to be the second most prevalent notifiable sexually transmitted infection in the US (CDC 2019a). The Gonococcal Isolate Surveillance Project (GISP), established by the CDC in 1986, has and continues to monitor trends in antimicrobial susceptibilities of *Ng* strains in the United States (Kirkcaldy 2016). Based on reported resistance of *Ng* strains, the CDC currently prescribes a two antibiotic protocol using ceftriaxone (a β-lactam) and azithromycin (a macrolide). Increased reports of these resistant strains have prompted an appeal by the CDC to researchers in the public and private sectors to intensify efforts to develop effective new treatments (CDC 2019b).

**Figure 1.**
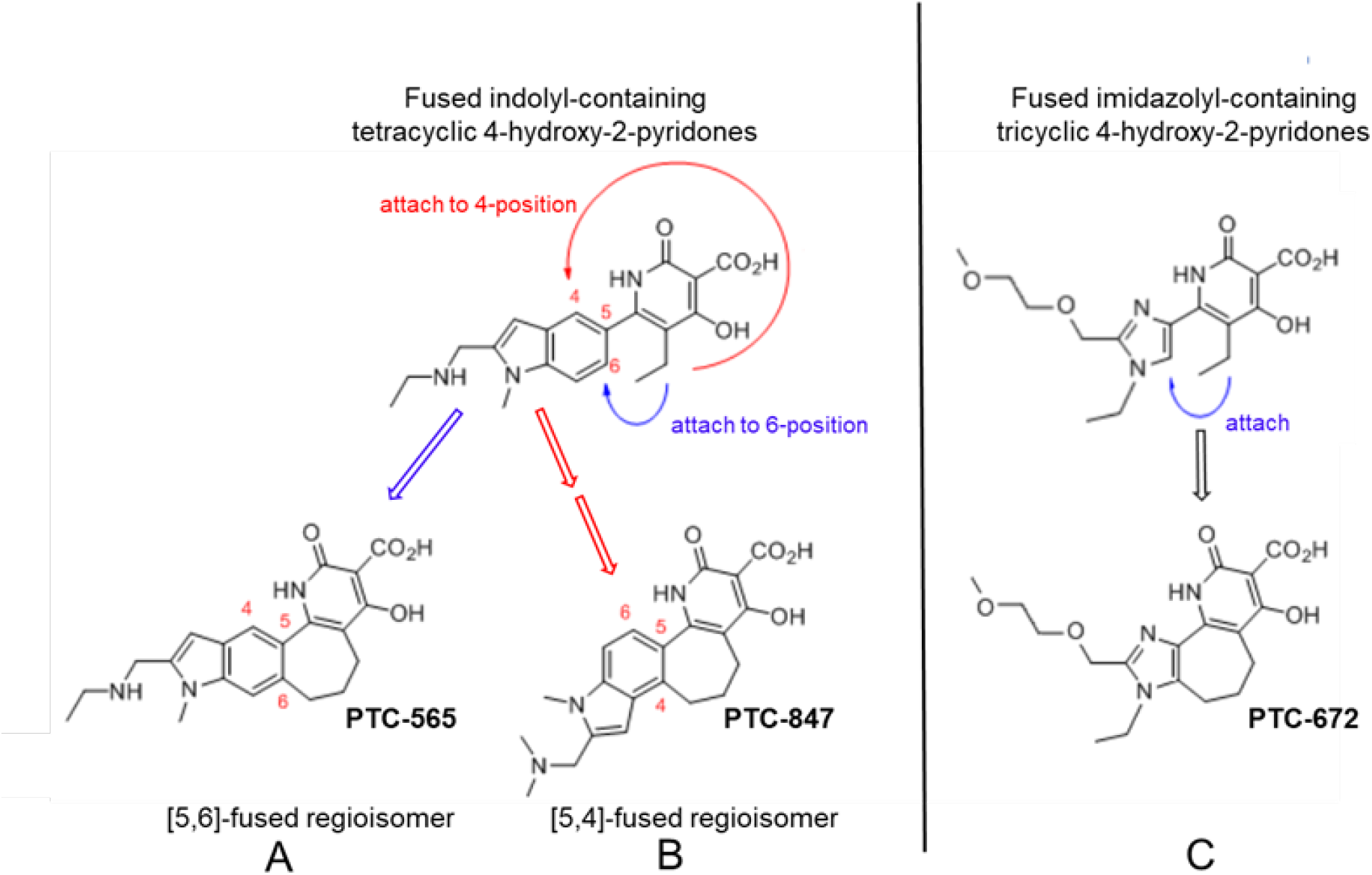
Structures of PTC-847 and PTC-672 with 4-hydroxypyridone nucleus. Three methylene units link the 4-hydroxy-2-pyridone core through the indole C-5 and C-4 position yielding two fused 7-membered ring regioisomers with restricted conformations, shown as compounds A and B. The [5,6]-fused regioisomer (PTC-565, A) is a broad spectrum DNA gyrase and topoisomerase IV inhibitor targeting gram negative pathogens (Gerasyuto 2018). The [5,4]-fused regioisomer with a basic dimethylamine group appended to the indole C-2 (PTC-847, B) is a potent class Ia RNR inhibitor selective for *Ng*. PTC-672 (C), a second selective *Ng* class Ia RNR inhibitor, has an imidazolyl moiety fused to the 4-hydroxy-2-pyridone through a 7-membered ring.

We now report that two new 4-hydroxy-2-pyridone pharmacophores (PTC-847 and PTC-672, Figure 1) inhibit many strains of *Ng* and MDR *Ng* by inhibiting biosynthesis of DNA. However, isolation of the *Ng* gyrase and topoisomerase IV reported herein revealed little inhibition by these compounds, in contrast to the related analogs in this class we previously reported (PTC-565, Figure 1A). In an effort to identify the target(s) of PTC-847 and PTC-672, we thus decided to select resistant *Ng* organisms to these compounds and to sequence their entire genomes.

Sequencing of several resistant strains revealed single nucleotide changes and amino acid substitutions, all located within the N-terminus of the large subunit (α) of the *Ng* class Ia RNR. RNRs catalyze the *de novo* conversion of the nucleoside diphosphates adenosine (A), guanosine (G), cytidine (C), and uridine (U) (collectively NDPs) to deoxynucleoside diphosphates (dNDPs) (Figure 2A). RNRs are essential in all organisms for DNA biosynthesis because of their role in controlling the relative ratios and amounts of dNTP pools. Imbalance in these pools increases mutation rates, replication and repair errors, and genome instability (Hofer 2012, Aye 2015, Greene 2020).

**Figure 2.**
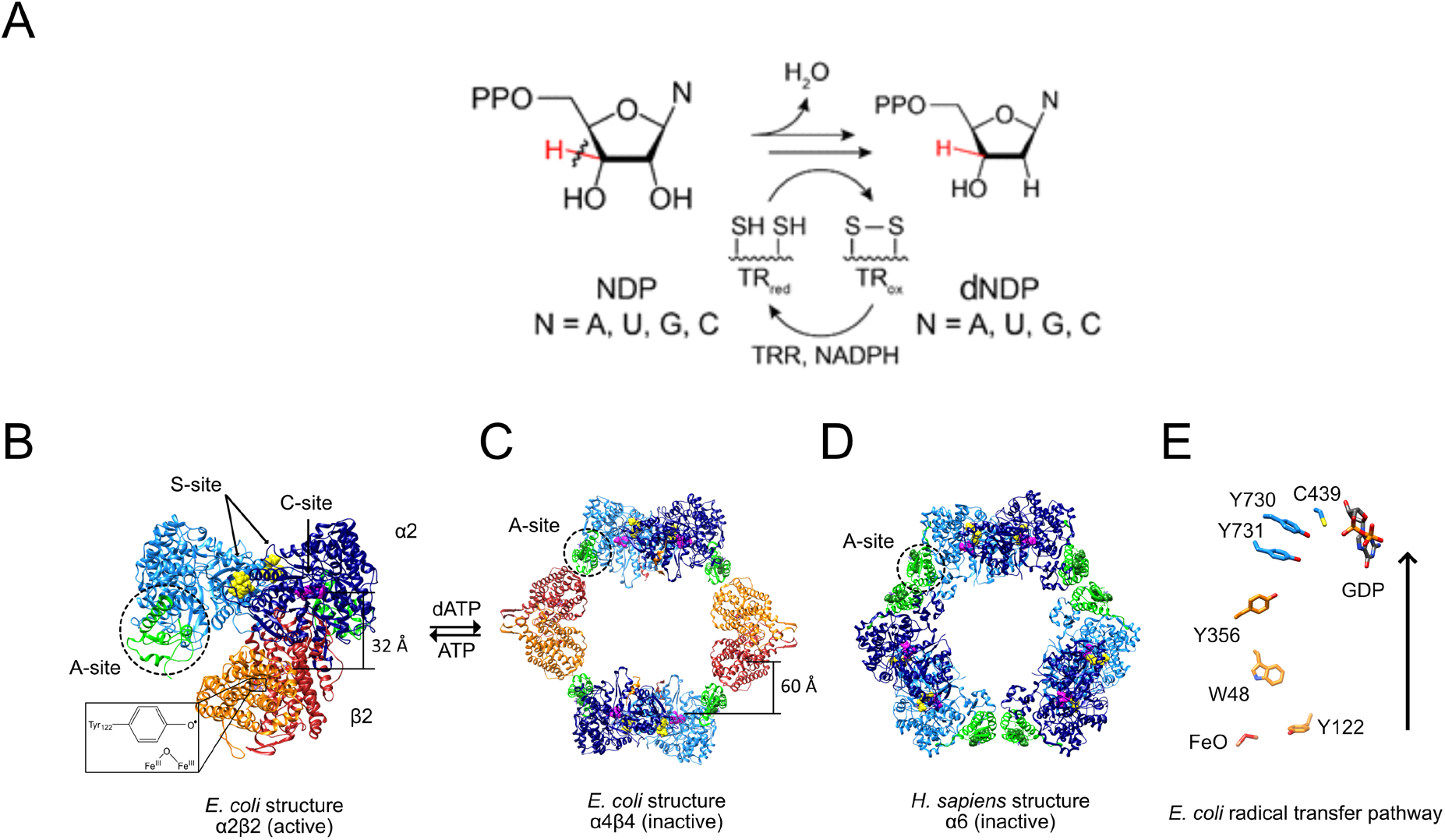
RNR reaction (A), structures of active and inactive states of Ia RNRs (B-D), and the radical transfer pathway (E). A. Each of the four NDPs are converted to their corresponding dNDPs, which involves cleavage of the 3′-C-H bond (red) and oxidation of two cysteines to a disulfide. Further turnover requires disulfide reduction by protein reductants thioredoxin (TR) and thioredoxin reductase (TRR). B. A cryo-EM structure of an active *Ec* RNR generated from a double mutant of β_2_ (F_3_Y122•/E52Q) incubated with α_2_, substrate GDP (C-site) and allosteric effector TTP (S-site) (Kang 2020). C. The inactive dATP-inhibited α_4_β_4_ state of *Ec* RNR. D. The inactive dATP inhibited α_6_ state of the human RNR. E. The essential diferric-tyrosyl radical cofactor and the components of the radical transfer pathway. A 32 Å distance is shown for the radical transfer pathway in B and 60 Å in C. In B-D, α2 is shown in blue with its N-terminal cone domain in green (A-site) surrounded by a dashed black circle. β_2_ is shown in red and orange.

Most organisms have multiple classes (designated class I, II and III) (Nordlund 2006, Greene 2020) of RNRs. They all share a common active site located in a structurally homologous protein (α, light and dark blue) and a common mechanism of NDP reduction. RNR classification is based on their essential and unique metallo-cofactors in β subunit that initiate the complex radical chemistry in α subunit (Eklund 2001, Norlund 2006, Cotruvo 2011). *Ng* possesses only one RNR, classified as a class Ia enzyme, that requires a diferric tyrosyl radical cofactor located in its second protein β (Figure 2B [orange, red], Eklund 2001, Kang 2020). Class I RNRs use α and β, two structurally homologous subunits, to form an active α_2_β_2_ complex (Figure 2B). The α subunit contains the catalytic site (C-site) and two allosteric effector sites, one involved in the regulation of substrate specificity (S-site) and the other in general enzymatic activity (A-site). The β subunit contains the metallo-cofactor required to initiate catalysis upon each NDP reduction event. What is most remarkable about the class Ia RNRs is that the metallo-cofactor in β must initiate complex reduction chemistry covering a 32 Å distance (Y_122_ β to C_439_ α) on each turnover (Figure 2B and E).

Class Ia RNRs, including the human enzyme, can form α_2_β_2_ active structures (Figure 2B) and are universally inhibited by dATP. The inhibited states, however, are distinct for the *Ec* and human class Ia RNRs (Figure 2C and D, respectively). The key to the structures of the inactive states is the *N*-terminal cone domain of subunit α (green, dotted circles in black). The dATP inhibited *Ec* RNR forms an α_4_β_4_ ring structure where the *N*-terminal cone domain that binds the dATP inhibitor interacts with β_2_. In the human dATP-inhibited state, the same *N*-terminal cone domain (green, dotted circles in black) plays a key role in formation of a trimer of α dimers in the hexameric α_6_ ring structure (Fairman 2011, Hofer 2001, Ando 2016, Greene 2020). Both inhibited states, by distinct mechanisms, prevent long-range radical initiation between the subunits and hence NDP reduction (Figure 2E). The mutations identified in the current studies are located in the *N*-terminal cone domain.

As introduced above, our present studies reveal that the 4-hydroxy-2-pyridone compounds, PTC-847 and PTC-672 (Figure 1), are potent inhibitors of class 1a RNRs, and demonstrate clear antibiotic activity preference toward all *Neisseria* species examined including MDR organisms and the *Nm* M2092 serogroup B (NMSB) reference strain. We also show that the compounds are not active against a panel of ten Gram-negative pathogens and commensal gut microorganisms. Evidence is presented for bactericidal activity against *Ng* associated with inhibition of DNA synthesis that does not result from inhibition of *Ng* purified DNA gyrase or topoisomerase IV, both fluoroquinolone targets. Selection for stable resistant *Ng* strains for each of these compounds and their subsequent whole genome sequencing led to the identification of mutations in the *N*-terminal cone domain of the *Ng* Ia RNR, suggesting the potential for inhibition of RNR by these compounds.

Using the structure of the *Ec* α_4_β_4_ inhibited state (Zimanyi 2016, Chen 2018), we show that the resistant mutants map to the *N*-terminal cone domain of the α subunit (green), which interacts with the β subunit (orange) of the α_4_β_4_ inhibited RNR structure (Figure 2C and Figure 3).

**Figure 3.**
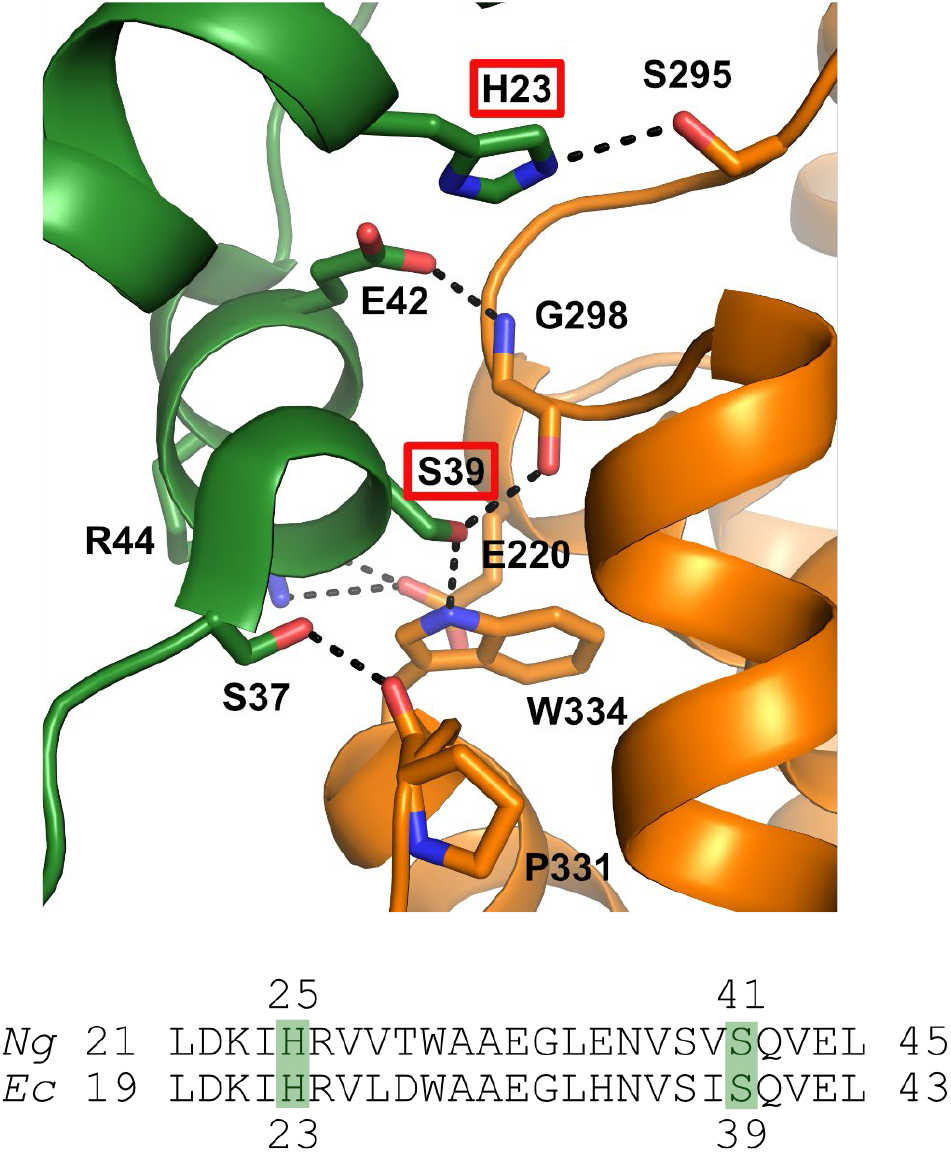
Location of mutations identified in the *Ng* Ia RNR by isolating resistant strains of*Ng* and mapping the mutation sites (H25R and S41L in α) where β is orange and α is green. The structure for the *Ec* class Ia RNR is shown with the residues in *Ec* (23, 39) that correspond to the mutation sites in *Ng* (residues 25 and 41) identified by red boxes (top) (Adapted from Chen 2018). Also shown is the sequence alignment of α in this region for *Ec* and *Ng*.

A high degree of sequence homology between the *Ng* and *Ec* class Ia RNRs suggested that the *Ng* enzyme might also form α_4_β_4_ structure. We report purification of the α and β subunits of the *Ng* class Ia RNR, characterization of its activity, and inhibition by the universal RNR inhibitor dATP and by PTC-847 and PTC-672. Using negative stain EM analysis, we demonstrate that *Ng* class Ia RNR can form a dATP-induced α_4_β_4_ state and that this state is abrogated by the mutations (H25R and S41L) that cause resistance to these compounds. These studies together suggest that the mode of class Ia RNR inhibition by the 4-hydroxy-2-pyridones might involve binding and potentiation of an inhibited α_4_β_4_ state.

To further characterize the potential use of these compounds as a treatment for infections caused by *Ng*, we assessed the efficacy of these compounds in both in vitro and in vivo mouse models. The compounds showed sustained clearance of infection when mice were infected vaginally with either antibiotic-susceptible or the MDR strains of *Ng*. The microbiological and biochemical studies together suggest that we have discovered novel compounds that selectively target the class Ia RNR for which no clinically useful inhibitors have previously been reported. These compounds also appear to inhibit class Ia RNRs by a new mode of action, which is altering the enzyme’s quaternary structure.

## RESULTS

### PTC-847 and PTC-672 are selective inhibitors against *N. gonorrhoeae* in culture

4-Hydroxy-2-pyridone analogs exemplified by PTC-847 and PTC-672 have been synthesized (Gerasyuto 2016, Wang 2016) to further explore the potential of this pharmacophore as a platform to uncover antibacterial agents targeting *Ng*. The minimum inhibitory concentrations (MIC) of PTC compounds were determined against six *Ng* strains (WHO strains, Figure 4A) that represent a full range of drug resistant phenotypes relevant to current WHO gonorrhea treatment guidelines (Unemo 2016), and ten Gram-negative pathogens and normally occurring gut organisms (Figure 4B). Two of these strains, WHO F (strain 13477) and WHO K (13479), were selected for a number of subsequent in vitro and in vivo experiments due to their respective sensitivity and resistance to a quinolone antibiotic, ciprofloxacin, that exhibits antibacterial activity by selectively inhibiting DNA synthesis. PTC-847 exhibited potent antibacterial activity against all six *Ng* strains with MICs ranging from 0.05 to 0.1 μg/mL. The weakly immunogenic NMSB strain had a MIC of 0.2 μg/mL (Table 1A). PTC-672 exhibited MICs ranging from 0.05 to 0.4 μg/mL against the six *Ng* strains, and a MIC of 0.1 μg/mL against the NMSB strain (Table 1A). The compound PTC-565 representing the [5,6] fused ring structure (Figure 1A) exhibited antibacterial activity against a broad spectrum of Gram-negative pathogens (Table 1 A and B); in contrast, PTC-847 and PTC-672 are inactive against the panel of Gram-negative pathogens and normal gut organisms evidenced by MICs ranging from 12.5 to ≥62.5 μg/mL (Figure 4B).

**Figure 4.**
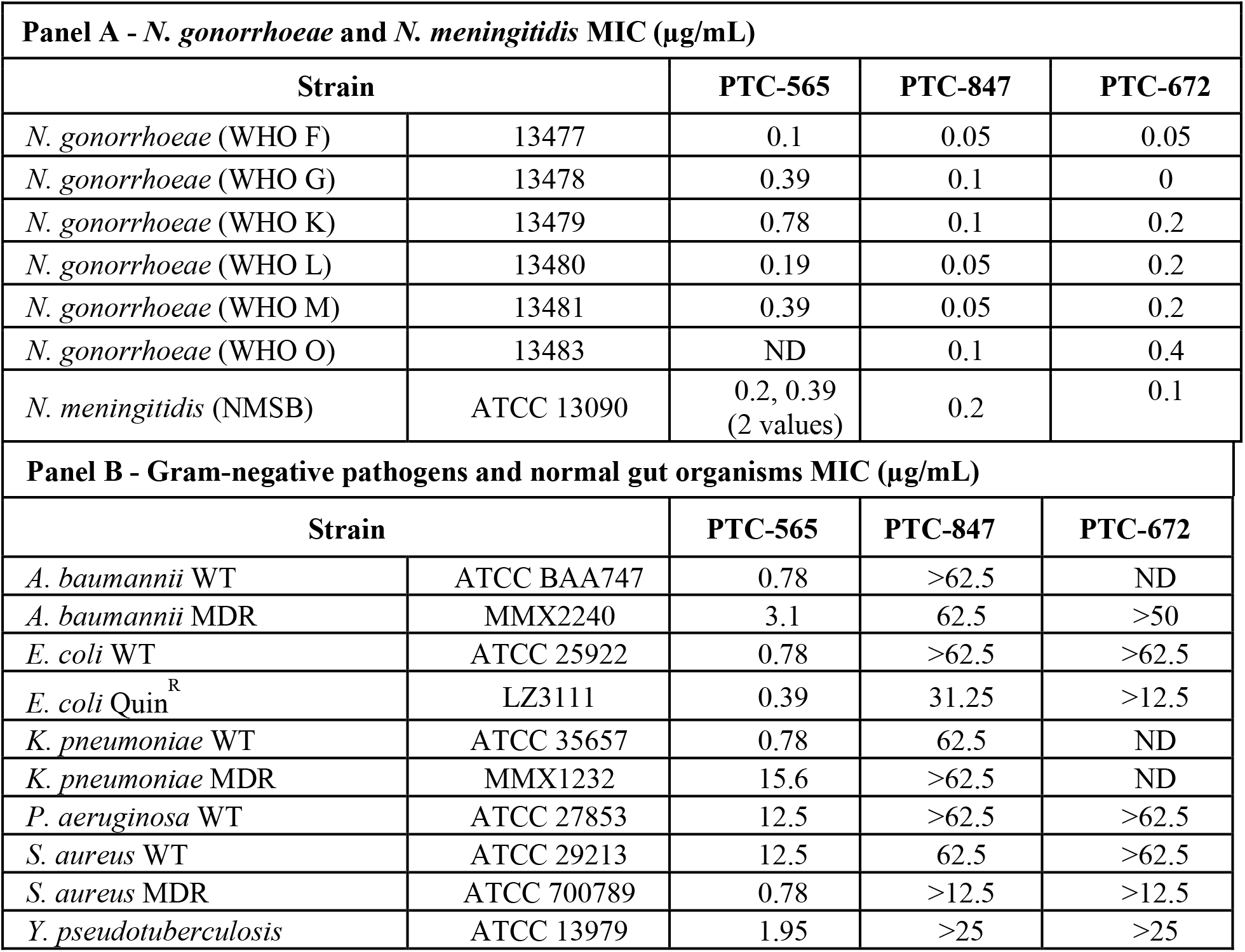
MIC of *Ng* strains (A) and Gram-negative pathogens and gut microorganisms (B). A) WHO F does not carry any antimicrobial resistance elements and is considered a susceptible strain. WHO G-M strains are resistant to quinolones. WHO K-O strains carry *penA*, *ponA*, *porB1*, and *mtrR* mutations associated with decreased susceptibility to cephalosporins. (Unemo 2016). *N. meningitidis* (NMSB) ATCC 13090 is the suggested reference strain for serogroup B (CLSI M22-A3 2016). B) This panel was constructed to test for inhibition of Gram-negative pathogens and normally occurring intestinal organisms, where WT = wild-type, MDR = multiple drug resistant, and Quin^R^ = quinolone-resistant strains. The *Nm*, Gram-negative, and normal gut organism strains were obtained from the American Type Culture Collection (ATCC), or were kindly provided by MicroMyx, LLC, (MMX), Kalamazoo, MI, or the laboratory of Lynn Zechiedrich (LZ), Baylor College of Medicine®, Houston, Texas. PTC-compound susceptibility testing was performed in accordance with the Clinical and Laboratory Standards Institute (CLSI) M07-A9 guideline (CLSI M07-A9 2012). Genus names: Acinetobacter (A), Klebsiella (K), Pseudomonas (P), Staphylococcus (S), Yersinia (Y).

The unique antibacterial activity against *Ng* strains with PTC-847 and PTC-672 (Figure 4) provided the impetus to examine them against a panel of 206 *Ng* strains with broad resistance phenotypes collected by the GIST Programme at Public Health England (Bolt 2016). Susceptibility testing resulted in MIC_90_ values for PTC-847 and PTC-672 equal to 0.12 μg/mL (Table S1). This collection includes strains lacking sensitivity to ceftriaxone and azithromycin, the current standard of care, and underscores that PTC-847 and PTC-672 are highly potent against *Ng*.

At concentrations of 1X MIC or greater, PTC-847 and PTC-672 exhibit time-dependent cidality against the *Ng* 13477 strain (Figure S1). In addition, a time-dependent post antibiotic effect was observed with PTC-847: following a brief exposure to and removal of PTC-847 where bacterial growth was persistently suppressed (Figure S2).

### PTC-847 targets DNA synthesis, but not DNA topoisomerases

Our previous studies (Gerasyuto 2018) established that a variety of [5,6]-fused indolyl-containing 4-hydroxy-2-pyridones (Figure 1A) targeted DNA synthesis, more specifically, fluoroquinolone resistant DNA topoisomerases, and inhibited a variety of gram-negative pathogens. To determine the target(s) of PTC-847, we followed the incorporation of radiolabeled precursors into total nucleic acids, DNA, and proteins in the *Ng* 13477 strain. The results revealed that inhibition of DNA but not protein synthesis appears to be responsible for growth inhibition by PTC-847 (Figure S3).

We then examined if inhibition of DNA topoisomerases (gyrase and topoisomerase IV) by PTC-847 was responsible for the observed DNA synthesis inhibition. Recombinant *Ng* 13477 gyrase (subunits A and B) and topoisomerase IV (both subunits) were cloned, expressed, and purified and the assay results compared to our previous similar studies on *Ec* DNA gyrase and topoisomerase IV. The PTC-847 half maximal inhibitory concentration (IC_50_) value for *Ng* gyrase was 19 μM and >32 μM for *Ng* topoisomerase IV and both *Ec* topoisomerases (Figure 5). Furthermore, PTC-847 at 100 μM did not appreciably inhibit the activity of human topoisomerase II. As a control, the fluoroquinolone antibiotic ciprofloxacin showed IC_50_ values ranging from 0.4 to 18 μM (Figure 5). These findings together suggest that topoisomerases are not the primary targets for PTC-847 in *Ng*.

**Figure 5.**
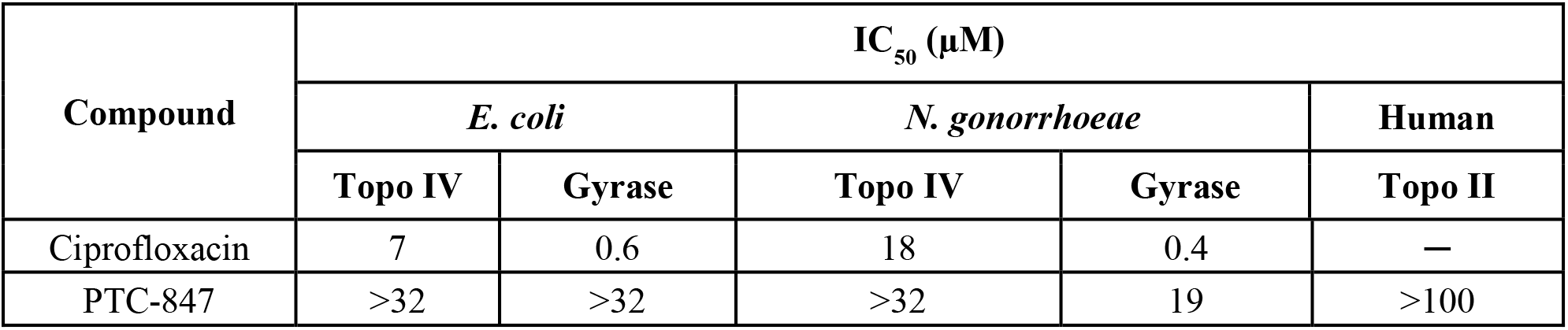
PTC-847 does not inhibit *Ng* gyrase or topoisomerase IV in vitro. Microplate-based supercoiling assays for DNA gyrase and topoisomerase IV were performed as previously described (Maxwell 2006). The sequence homology between the *Ng* and *Ec* enzymes are 70% and 75% in gyrase subunit A and B, respectively, and 64% and 65% in topoisomerase IV subunits A and B, respectively. The assays thus followed the *Ec* protocols and are described in detail for the *Ng* enzymes in the Methods. Decatenation assays using purified gyrase AB and topoisomerase IV proteins were performed following the protocols for Profoldin’s topoisomerase II and IV DNA decatenation assays (ProFoldin, Hudson, MA) also in Methods.

### PTC-847 targets the class Ia RNR large subunit

In an effort to identify the target of inhibition of PTC-847, we used *Ng* 13477 (Figure 4) to select and isolate a stable PTC-847-resistant isolate (PTC-847^R^) (Table S2). The frequency of resistance for PTC-847^R^ was on the order of 10^−9^ which is similar to that observed for resistance to ciprofloxacin or ceftriaxone. PTC-847^R^ was tested against a wide variety of antibiotics having different modes of action (Table S3). The PTC-847^R^ strain was as sensitive to all classes of antibiotics as the wild-type *Ng* 13477 strain, except for susceptibility to PTC-847. The MIC for PTC-847 against *Ng* 13477 was 0.05 μg/mL compared to 15.6 μg/mL for the PTC-847^R^ isolate.

To identify the specific PTC-847 molecular target(s), the PTC-847^R^ isolate was subjected to whole genome sequencing using the *Ng* 13477 WT strain as the reference. A single C to T transition that results in a serine (S) to leucine (L) amino acid substitution at position 41 (S41L) in the α-subunit of the *Ng* class Ia RNR was identified (Figure 3). The gene sequence change was verified by comparing DNA sequences from two of the resistant PTC-847^R^ clones against the WT reference strain sequence. The target site conferring resistance to PTC-847 was further confirmed in an isogenic strain (designated PTC-847^S41L^) made by transforming the WT 13477 strain with a DNA fragment containing the α-subunit gene mutation, followed by selection on agar plates containing PTC-847. Hundreds of PTC-847 resistant colonies were obtained when the 13477 strain was transformed with the mutated, but not the WT, α-subunit gene sequence.

A similar strategy using PTC-672 resulted in a stable resistant strain PTC-672^R^. Sequencing the PTC-672^R^ class Ia RNR gene revealed a single A to G transition that resulted in a histidine (H) to arginine (R) amino acid substitution at position 25 (H25R) in the α-subunit (Figure 3). Thus, both inhibitors target the large subunit of the *Ng* Ia RNR that together with its small subunit are essential for DNA synthesis by catalyzing the conversion of NDPs to dNDPs. In support of the mechanism of inhibition of PTC-847, an increased ratio of NTPs to dNTPs in the susceptible WT 13477 strain was observed subsequent to treatment with the inhibitor at 1x MIC for one hour followed by nucleotide extraction and LC/MS analysis (Chen 2009) (Table S4).

### Isolation and activity characterization of *Ng* Ia RNR

The active form of the class Ia RNR is α_2_β_2_ (Figure 2B). The two subunits, however, have weak affinity with the K_D_ ~0.2 μM for *Ec* and human RNRs; therefore, each subunit was cloned, expressed, and purified independently (Greene 2020). Furthermore, assay protocols are optimized for each RNR from each organism. For *Ng* α, an expressed tagged α subunit protein was purified by affinity chromatography and the tag removed to eliminate potential interference with the α_4_β_4_ quaternary structure (Figure 2B and 2C) and enzyme activity. Subunit β was cloned and expressed with an N-terminal (His)_6_-tag and purified to homogeneity to give 0.7 to 0.9 diferric tyrosyl radical cofactors (Figure 2A, Methods, Figure S4). The activity of *Ng* RNR was optimized (Figure S4) so that its inhibition by dATP and PTC-847 and PTC-672 could be determined. The activity at 0.1 μM and 1:1(α:β) ratio of subunits was 1300 nmol min^−1^ mg^−1^ with GDP/TTP as substrate and effector, similar to that for *Ec* RNR (2500 nmol min^−1^ mg^−1^ with CDP/ATP as substrate and effector) (Methods, Figure S4). The apparent *K*_*m*_ is 0.03 μM for *Ng* RNR and as noted above that for *Ec* and human RNRs is ~0.2 μM. The lower *K*_*m*_ and the basis for the drop off in activity at > 0.3 μM (Figure S4) require further investigation.

All class Ia RNRs have N-terminal cone domains of α (~100 amino acids) that control enzyme activity (A-site Figure 2A, ATP activates and dATP inhibits). Studies of *Ng* RNR activity as a function of dATP concentration (Figure S5) reveal a profile very similar to *Ec* RNR, including biphasic kinetics based on dATP binding to both the S-site and A-site (Hofer 2011). This result is expected given the 76% sequence identity for α between *Ng* and *Ec*, including the H25 and S41 (Figure 3) found altered in *Ng* resistant to the PTC compounds. Human α, on the other hand, shares only 30% sequence identity to *Ng* and does not have residues homologous to either H25 or S41. Inhibition studies with H25R and S41L *Ng α* mutants showed much lower sensitivity to dATP (Figure S5).

The *Ng* GDP/TTP (Figure S4) and [^3^H]-CDP/ATP (Figure S5) assays were used to assess the ability of PTC-847 and PTC-672 to inhibit *Ng* RNR activity. The results are summarized in Figure 6 and Figure S6. PTC-847 exhibited 93% inhibition at 4 μM PTC-847, whereas PTC-672 showed 78% inhibition at 2.5 μM. Similar experiments with *Ec* and human RNRs using the [^3^H]-CDP/ATP assay were also carried out. As expected, based on sequence similarity, the *Ec* RNR is also inhibited, whereas the human enzyme, even at 100 μM inhibitor concentration, still retains substantial activity (Figure 6).

**Figure 6.**
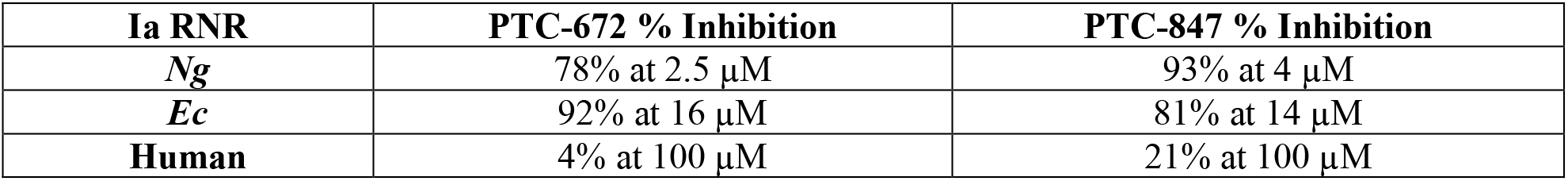
Inhibition of class Ia RNRs in vitro. Various concentrations of PTC-672 or PTC-847, or 100 mM N_3_CDP (positive control) were added to 37 °C reaction mixtures containing α, β, ATP, TR, TRR, and NADPH. The mixtures were incubated for either 30 sec, 5 min or 15 min prior to initiation of the reaction with 5-[^3^H]-CDP. Aliquots were quenched at 1, 2, 3 and 4 min in 2% HClO_4_. All samples were neutralized by the addition of 0.5 M KOH and processed following a standard protocol (Ravichandran 2020). Percent inhibition was calculated relative to a 1% DMSO negative control.

### Negative stain EM reveals *Ng* Ia RNR can form α_4_β_4_ while mutations resulting in PTC-847^R^ and PTC-672^R^ cannot

We have previously shown using negative stain EM analysis with *Ec* RNR that dATP shifts active RNR into the inhibited α_4_β_4_ state (Figure 2C), which is readily observed due to the ring structures (Ando 2011). Comparable experiments with *Ng* Ia RNR were carried out under very similar conditions to the inhibition studies and the results are shown in Figure 7. Panel A shows that, in the presence of 1 mM dATP, ring structures are present. The 2D classifications and 3D reconstruction are shown in Figure 7B and C, and reveal that *Ng* Ia RNR, like the *Ec* enzyme, forms the α_4_β_4_ ring structure inhibited state induced by dATP.

**Figure 7.**
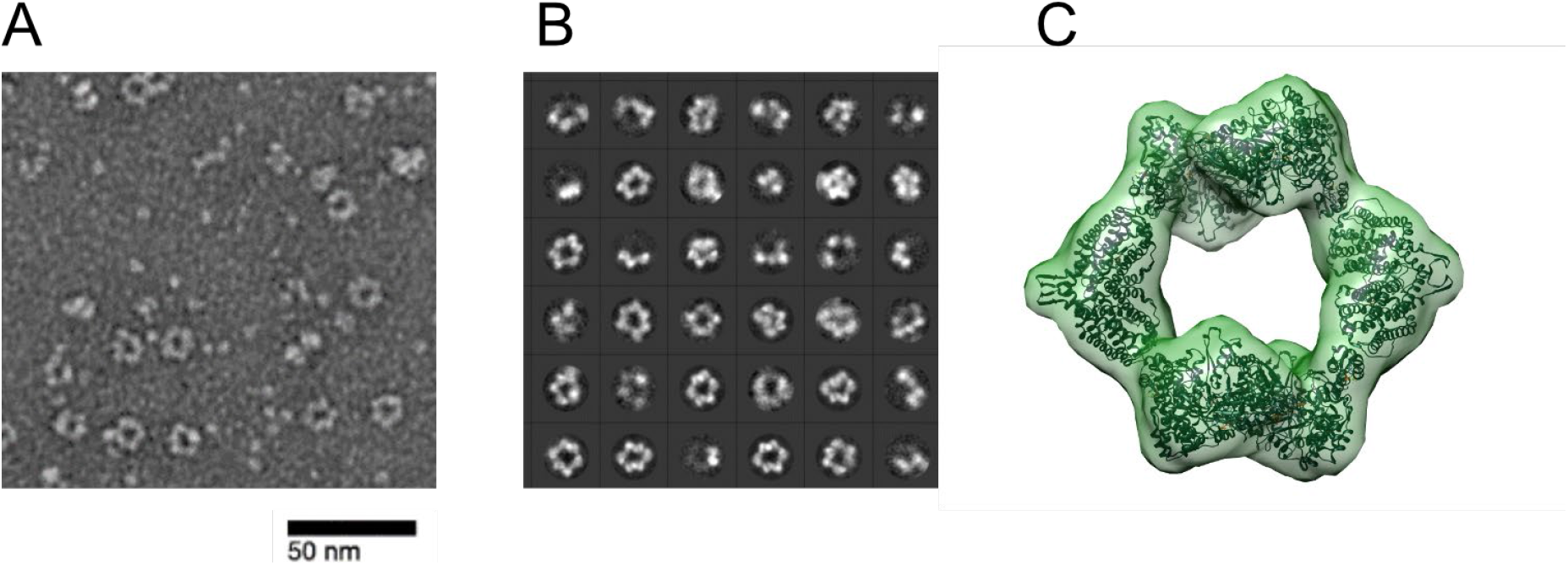
N. gonorrhoeae RNR forms rings under inactivating conditions observed with Ec RNR. (A) Representative negative stain image taken at 84000x nominal magnification (1.79 Å/pixel) of 1:1.5 α:β (15 ng/μL final concentration; 0.1 μM α_2_ and 0.15 μM β_2_) in the presence of 1 mM dATP and 1 mM CDP. Sample was prepared at 4 °C before staining. (B) Representative 2D class averages (36 shown of 200 total) from 27,109 picked particles from grids prepared under the same conditions as (A). (C) Negative stain reconstruction at 21 Å resolution of *Ng* class Ia RNR α_4_β_4_ with *Ec* class Ia RNR α_4_β_4_ crystal structure fit to the density (PDB 5CNS; Zimanyi et al. 2016).

Our hypothesis for RNR inhibition by the 4-hydroxy-2-pyridone derivatives, based on the mapped resistance mutations (Figure 3 and Figure S5), is that they potentiate α_4_β_4_ formation. To determine if the selected resistant mutants affected the extent of α_4_β_4_ formation, PTC-672^H25R^ and PTC-847^S41L^ α subunits were prepared and purified by the same methods as for the WT proteins and examined as described in Figure 7. As shown in Figure 8, in the WT control, α_4_β_4_ ring structures are clearly visible whereas no such structures are observed with either mutant. An additional control in which ATP (an activator) replaces the dATP inhibitor, also reveals no α_4_β_4_ structures for either the WT or mutant proteins (data not shown).

**Figure 8.**
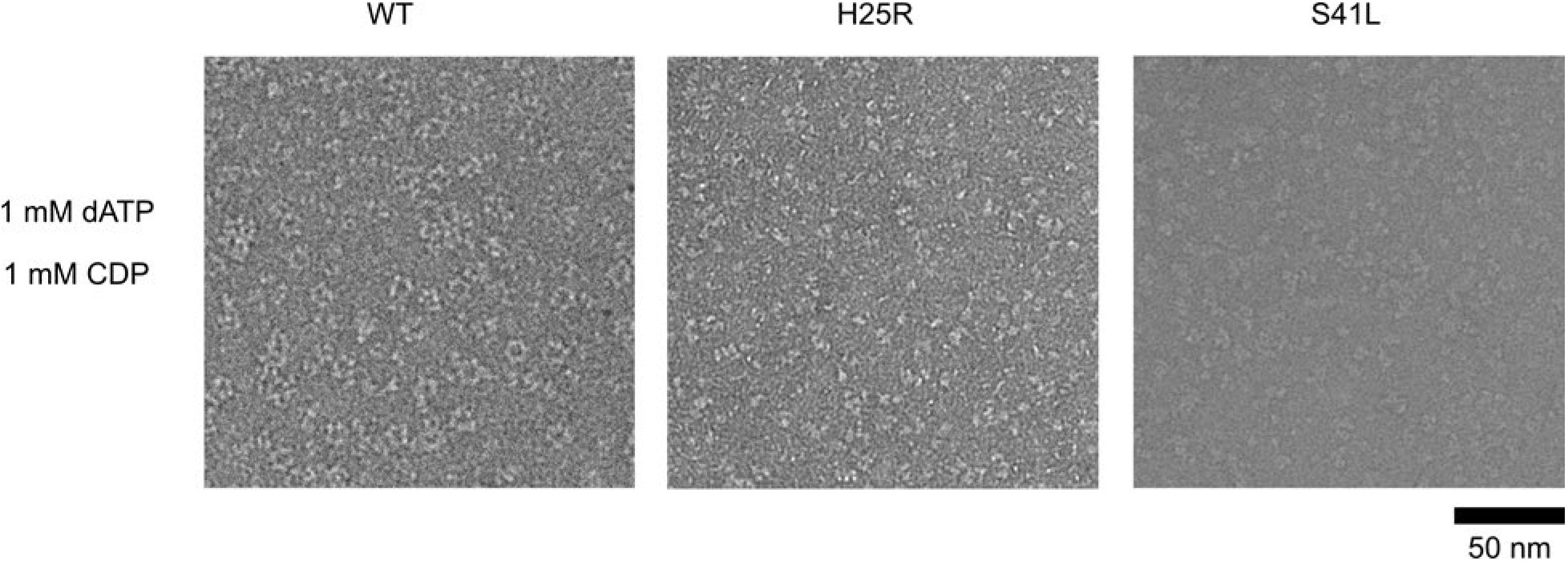
Ng RNR S41L and H25R variants do not form rings under dATP inhibiting conditions. WT, PTC-672^H25R^, or PTC-847^S41L^ α subunits were combined with WT β in a 1:1.5 ratio (15 ng/μL final concentration; 0.1 μM α and 0.15 μM β) in the presence of 1 mM dATP and 1 mM CDP and imaged on an F30 (FEI) microscope. Samples were prepared at 4 °C prior to staining.

The results of these studies support our model that inhibition by PTC-847 and PTC-672 require an α_4_β_4_ state and that they potentiate α_4_β_4_ formation. The dependency on a dATP-induced inactive α_4_β_4_ state also explains why the *Ng* and *Ec* Ia RNRs are both sensitive to these inhibitors, whereas the structurally distinct human Ia RNR α_6_ inhibited state is not. This result provides the first experimental support for the idea that an inhibited state of a bacterial RNR might be a new way to selectively target the bacterial RNRs.

### Assessment of PTC-672 and PTC-847 for treatment of *Ng* infections

In the following sections an in vivo mouse model for vaginal infection with drug susceptible and resistant strains of *Ng* was used to monitor infection clearance. *Ng* 13477 and 13479 (WHO F and K, “susceptible and resistant strains to PTC-672, see Table 1) were constructed for in vitro and in vivo fitness testing. These strains were used to monitor the development of resistance to the test compounds in vivo and to access bacterial survival.

### In vivo studies of *Ng* infections using a mouse model

*Ng* has adapted for the human vaginal tract and will not grow in other species. With administration of high doses of estrogen to ovariectomized female mice and depletion of the normal vaginal bacterial flora, however, mice can be infected with *Ng.* (Raterman 2019). To better control the levels of estrogen, these mice are administered estrogen by slow release pellet two days prior to infection. Concomitantly, they are administered vancomycin, streptomycin, and trimethoprim to deplete commensal flora. These studies are accompanied by administration of streptomycin (Strep) once daily for the remainder of the studies to out-compete the normal flora (for details see Methods).

### *Ng* strain creation for efficacy and fitness studies

As a starting point for susceptibility testing with PTC-672, the *Ng* 13477 strain, which lacks antimicrobial resistance elements (Unemo 2016), was tested against the MDR *Ng* 13479 strain that has a high level of resistance to quinolones and carries mutations associated with decreased susceptibility to cephalosporins (Unemo 2016). Four new isogenic strains were created for the in vivo studies from *Ng* 13477 and 13479. One set was engineered to have PTC resistance (PTC-672^H25R^) and the second to have resistance to both PTC-672 and streptomycin (Strep^R^-PTC-672^H25R^). They are designated Iso 13477 Strep^R^-PTC-672^H25R^ and Iso 13479 Strep^R^-PTC-672^H25R^. The strains were created using a PCR product containing the H25R α-subunit gene and/or DNA isolated from the streptomycin resistant *Ng* FA1090 laboratory strain.

### In vitro selectivity of PTC-847 and PTC-672

To test if normal intestinal flora are resistant to inhibition by these compounds, PTC-847 and PTC-672 MIC values were determined against a panel of normally occurring intestinal organisms (Thursby 2017). PTC-847 and PTC-672 spared the gut flora relative to the comparator solithromycin (Table S5), a fourth-generation macrolide with enhanced activity against macrolide-resistant bacteria (Farrell 2016). Furthermore, PTC-672’s effect on several commensal *Neisseria* strains was assessed and it was determined that PTC-672 inhibited their growth to a lesser extent than that of *Ng* (Table S6). Based on the demonstrated target and biological selectivity of PTC-847 and PTC-672, these compounds may be better tolerated compared with current therapies.

### In vivo efficacy of PTC-672 using the mouse model with susceptible and MDR *Ng* strains

The efficacy of PTC-672 was assessed in the mouse model. On Day 0 of the study, mice were infected vaginally with the *Ng* Iso 13477 Strep^R^ or Iso 13479 Strep^R^ strains. Beginning on Day 1, mice were swabbed daily for one week to determine the bacterial load. On Day 2, mice were randomized into groups based on Day 1 bacterial load and administered vehicle, ciprofloxacin, ceftriaxone, or PTC-672. Efficacy is defined as complete and sustained clearance of infection at Day 5 post-treatment.

In mice infected with the Iso 13477 Strep^R^ strain (Figure 9A), those that received vehicle had a robust infection for 7 days, with 5/9 mice remaining infected through Day 7. Treatment with a single oral dose of ciprofloxacin at 30 mg/kg resulted in complete clearance of the infection in 10/10 mice (~4-log drop in bacterial infection) within 24 h following treatment. PTC-672 administered as a single oral dose (10, 15, 20, 25 or 30 mg/kg) resulted in clearance of infection at all doses in 10/10 mice within 24 h. The clearance of infection by ciprofloxacin or PTC-672 was sustained through Day 7.

**Figure 9.**
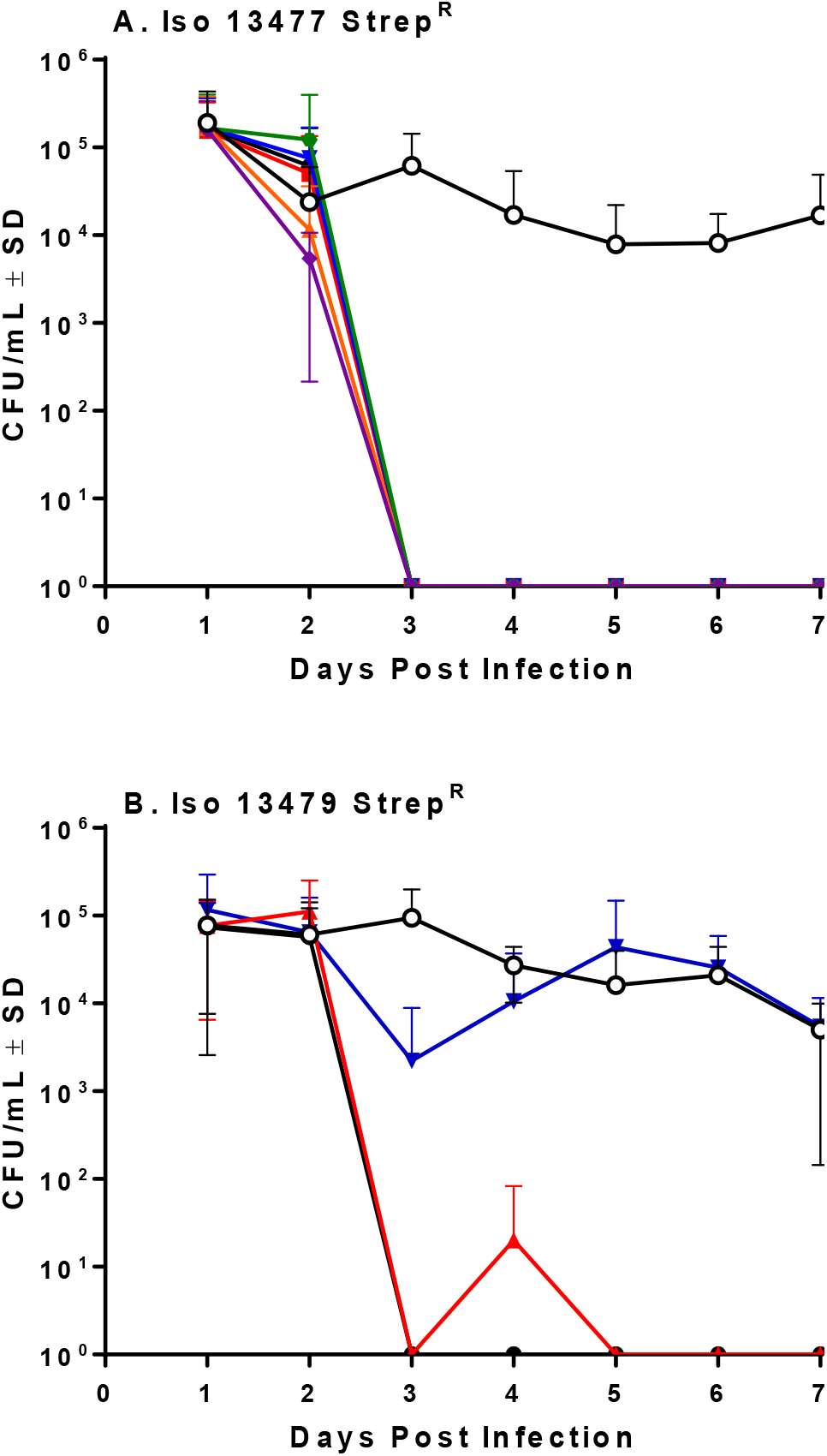
PTC-672 in vivo efficacy in a mouse model of gonorrhea. Mice were vaginally inoculated with streptomycin resistant *Ng* strains and swabbed daily over a one-week period. At each sampling time point, the number of bacteria (CFU/mL) were counted by plating onto selective plates and the mean CFU/mL ±SD values were determined. The mean ±SD values were plotted. A) Symbols: vehicle open black, ciprofloxacin solid black, PTC-672 10 mg/kg purple, 15 mg/kg green, 20 mg/kg orange, 25 mg/kg blue, and 30 mg/kg red. B) Symbols: vehicle open black, ceftriaxone solid black, PTC-672 30 mg/kg blue and 60 mg/kg red.

In mice infected with the Iso 13479 Strep^R^ strain (Figure 9B), those that received vehicle had a robust infection for 7 days of study, with 8/10 mice remaining infected through Day 7 of study. Treatment with a single intraperitoneal dose of ceftriaxone (100 mg/kg) resulted in complete clearance of the infection in 10/10 mice within 24 h and clearance of infection was sustained until Day 7. PTC-672 administered as a single oral dose (60 mg/kg) resulted in complete clearance of the infection in 10/10 mice within 24 h of dosing and was also sustained until Day 7. At the lower dose of 30 mg/kg PTC-672 resulted in an average 1.46-log drop in bacterial load (8/9 mice cleared infection) 24 h post-dose. However, infection in several mice re-bounded on subsequent days of the study suggesting that a dose of 30 mg/kg was sub-efficacious in the Iso 13479 Strep^R^ model.

### In vitro fitness testing of PTC-672^R^

The relative fitness cost associated with PTC-672 resistance was determined in vitro by the competition strains described above. Competitive fitness assays measure net growth of the two populations of isogenic resistant and susceptible strains and account for differences between these populations across the full growth cycle including lag times, exponential growth rates, and stationary phase dynamics.

In this in vitro study, Iso 13477 Strep^R^-PTC-672^H25R^ or Iso 13479 Strep^R^-PTC-672^H25R^ and Iso 13477 Strep^R^ or Iso 13479 Strep^R^ were mixed, inoculated into growth media, and aliquots taken at intervals over a 24 h incubation period. At each sampling time point, the number of total (susceptible + resistant) bacteria and resistant bacteria were determined by plating onto non-selective and PTC-672-supplemented plates, respectively.

Relative fitness is expressed as the competitive index (CI), the ratio of bacterial burdens between the PTC-672 resistant and susceptible strains at each time point divided by the baseline ratio at the beginning of the experiment. CI of ~1 indicates that the fitness of the resistant bacteria is similar to that of the parent bacteria, CI >1 indicates that the resistant bacteria are more fit, and CI <1 suggests that the resistant bacteria are less fit in the absence of selection pressure. In this experiment, the CI for *Ng* Iso 13477 Strep^R^-PTC-672^H25R^ and Iso 13479 Strep^R^-PTC-672^H25R^ decreased exponentially to a CI of 0.06 at 8h (Figure 10), demonstrating that the resistant bacteria are less fit.

**Figure 10.**
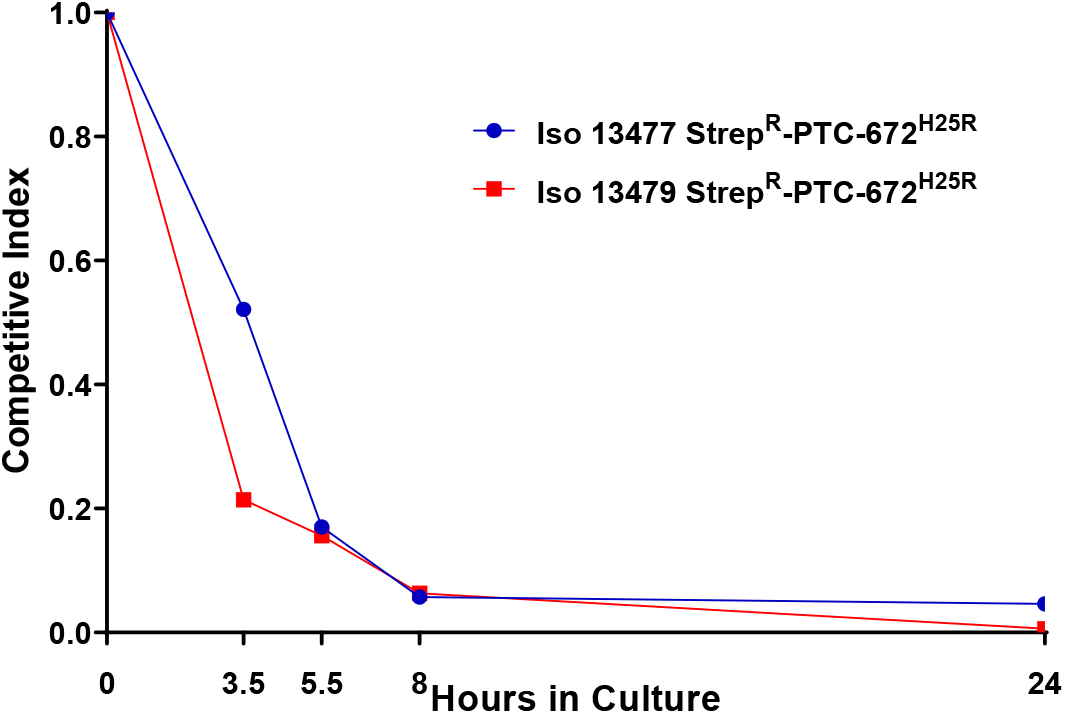
H25R mutation in *Ng* RNR reduces fitness in vitro. Iso 13479 Strep^R^-PTC-672^H25R^ (red) and Iso 13477 Strep^R^-PTC-672^H25R^ (blue) were mixed in a 1:1 ratio, inoculated into fresh growth media, and cultured. At each sampling time point, the numbers of total and resistant bacteria were determined by plating onto non-selective and PTC-672-supplemented plates, respectively. Relative fitness was expressed as the CI (see text).

### In vivo fitness testing of PTC-672 resistant *Ng*

For these experiments, mice pre-treated as described above were inoculated vaginally with either the isogenic Strep^R^-PTC672^H25R^ strain or a mixture of the Strep^R^ parent and Strep^R^-PTC672^H25R^ strains and swabbed daily over a one-week period. At each sampling time point, the relative fitness expressed as the CI was determined by plating onto non-selective and PTC-672-supplemented plates.

When infected with only the Iso 13477 Strep^R^-PTC-672^H25R^ or the Iso 13479 Strep^R^-PTC-672^H25R^ strain, the infection was similar to that elicited with the Strep^R^ parent strain. A similar number of mice were infected on Day 1 and the infection level was comparable throughout the study. For fitness testing with mixed strains, the mean and median CIs for *Ng* Iso Strep^R^-PTC-672^H25R^ 13477 and Iso 13479 Strep^R^-PTC-672^H25R^ resistant strains decreased to <0.01 by Day 2 post infection (Figure 11).

Thus, although the PTC-672 resistant *Ng* strains infection levels in mice were comparable to those for the isogenic PTC-672 susceptible parent strains, when the two isogenic strains were inoculated in the same anatomical location in vivo, the PTC-672 resistant isolates did not successfully compete with the parent strains. These data suggest that if PTC-672 resistance does arise in *Ng* strains in situ, the PTC-672 resistant strain would be less fit and thus less likely to propagate. These results combined with the outcomes of the in vitro fitness study suggest that PTC-672 may lessen the emergence of resistance compared with current therapies.

**Figure 11.**
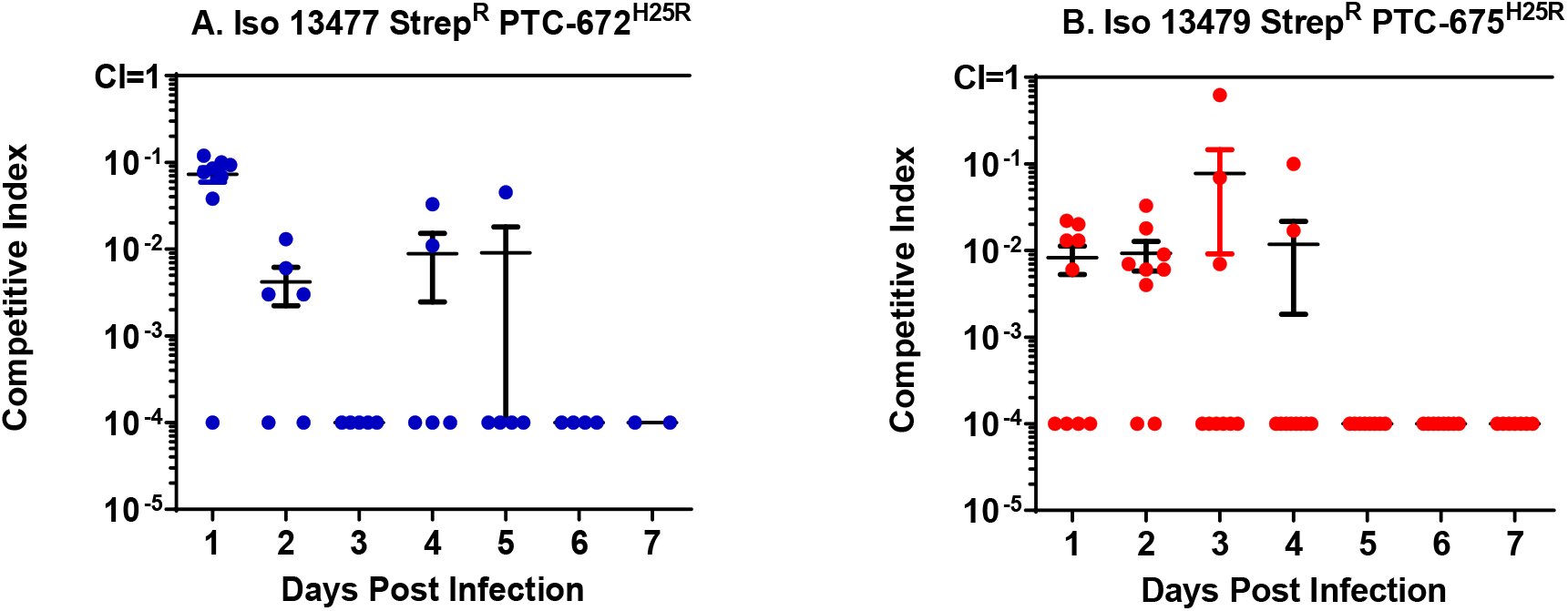
H25R mutation in *Ng* α of its Ia RNR reduces fitness in vivo. Mice were vaginally inoculated with the mixed isogenic resistant and susceptible strains and swabbed daily over a one-week period. The CI was determined at each sampling time point as described above. The medians ±ranges are shown as solid horizontal lines.

## DISCUSSION

Novel strategies for developing treatments for highly resistant bacterial pathogens are needed. As a result, there is increased interest in developing single-pathogen therapies against multidrug-resistant organisms. The discovery that the 4-hydroxy-2-pyridone compounds with 5,6 fused ring structures (Gerasayuto 2016) exhibited broad spectrum bactericidal activity against many gram negative pathogens by targeting topoisomerases and inhibiting DNA synthesis provided the impetus for further investigation of a general pharmacophore. In conjunction with the reports of increasing MDR in *Ng* by the WHO and CDC, this organism became the focus of our efforts. The discovery that the 4,5-fused pyridines with indolyl and imidazolyl moiety rings (Figure 1B, C) were potent and selective inhibitors of *Ng, Nm,* and MDR *Ng* (Figure 4) led to further studies to identify the target(s) of these compounds and the suitability of two compounds (PTC-847 and PTC-672) as potential single organism therapeutics.

We demonstrated that these compounds, although inhibiting DNA synthesis, do not appear to target *Ng* gyrase or topoisomerase IV, but rather uniquely target *Ng* Ia RNR. The dramatic increase in life threatening bacterial infections, coupled to the increase in resistance and MDR to widely used therapeutics (Walsh 2015) makes identification of this new therapeutic target particularly exciting.

This discovery was made through mapping mutants resistant to these compounds to the N-terminal cone domain (100 amino acids) of the *Ec* Ia RNR (Figure 2B, 2C). This domain forms the allosteric activity site responsible for the dATP-induced, inhibited α_4_β_4_ state (Figure 2D). We have previously shown using negative stain EM analysis with *Ec* Ia RNR that dATP shifts the active α_2_β_2_ enzyme into an inhibited α_4_β_4_ state, readily observed by its distinct ring structure (Ando 2011). Comparable experiments now show that *Ng* Ia RNR enzyme also forms an inhibited ring structure when induced by dATP (Figure 7). Studies in vitro (Chen 2018) and in vivo (Ahluwalia 2012) of *Ec* RNR establish that α/β interface residues H23 and S39 (H25 and S41 in *Ng* RNR) disrupt this quaternary structure and our preliminary data obtained using EM suggest the same is true for *Ng* RNR (Figure 7). The detailed mechanism by which dATP and PTC compounds inhibit the *Ng* RNR are currently being investigated as many questions about the wild-type and mutant αs and their affinity for β_2_, and the stoichiometry for dATP, PTC-847 and −672 remain unanswered.

Trapping distinct quaternary structures of regulatory proteins with small molecule inhibitors has been successfully exploited with identification of the importance of inhibited protein kinase states in targeting signaling pathways involved in the regulation of cancer cell growth (Greene 2020). Recently, biochemical and cell-culture studies have revealed that clorfarabine (ClF), a nucleoside therapeutic used in the treatment of acute lymphoblastic and acute myeloid leukemias, subsequent to its phosphorylation to a di- or trinucleotide (ClFDP, ClFTP) binds to the α subunit of human Ia RNR and potentiates formation of the α_6_ inhibited state (Aye 2011, Brignole 2018). In contrast with the dATP-induced α_6_ inhibited state (Fairman 2011), which rapidly reverts to a mixture of α monomers and dimers on dATP dissociation, both the ClFDP and ClFTP α_6_ states remain hexameric for some time subsequent to their dissociation (Brignole 2018, Aye 2011). Interestingly the *E. coli* Ia RNR, which does not form an α_6_ state, is not inhibited by ClFTP. While the unique bioinorganic and organic chemistry of RNRs have resulted in targeted inhibition of the active site, α/β subunit interactions and formation of the active metallo-cofactor (Cotruvo 2011, Greene 2020), the unusual stability of the ClFDP or ClFTP-induced α_6_ state suggests a novel strategy for selectively targeting this essential enzyme in other organisms.

Despite the structural homology between *Ng* and human α, no appreciable inhibition of the human RNR by PTC-847 or PTC-672 is observed (Figure 6), consistent with the distinct α_6_ quaternary structure. The discovery that PTC-847 and PTC-672 inhibited *Ng* but not human Ia RNR suggests that trapping the inhibited quaternary state might be a new approach for therapeutics that selectively target *Ng*.

A number of observations with PTC-847 and −672 further support their potential as new therapeutics. PTC-847 and PTC-672 inhibit all strains of *Ng* and *Nm* (Table 1), but examination of a panel of Gram-negative pathogens and normally occurring intestinal flora including commensal *Ng* strains were found to be insensitive (Figure 4, Table S5, and Table S6). We postulate that this selectivity may be explained by the observation that *Ng* and *Nm* possess only a single RNR, whereas many microorganisms have multiple RNRs (additional class Is [Ib-Ie] or class II and III) (Lundin 2009, Stubbe 2019) that may not be targeted by these compounds. Based on the demonstrated target and biological selectivity of PTC-847 and PTC-672, these compounds may be better tolerated compared with current therapies.

We have demonstrated in vivo efficacy of PTC-672 in a mouse model of *N. gonorrhoeae*. When administered as a single oral dose of 60 mg/kg PTC-672, complete clearance of infections was observed with both a susceptible and an MDR strain of *Ng* in 10/10 mice within 24 h following dosing. We also showed that the H25R–α mutation reduced the fitness of the susceptible and MDR *Ng* strains in vitro and in vivo competition experiments. In vivo we showed effectiveness of PTC-672 and PTC-847 in quinolone resistant and WT *Ng* strains and that if resistant isolates arise, they are less fit and will not likely become dominant. Taken together, these results suggest that treatment with PTC-847 or PTC-672 may be more selective, better tolerated, and lessen the emergence of resistance compared with current therapies.

Resistance of life-threating bacterial infections to most types of antibiotics has been globally emerging over recent years due in part to overprescribing. One possible strategy is to restrict broad spectrum antibiotic use to treatment of serious MDR infections and to develop novel single-pathogen agents for treating low prevalence infections. However, there is some concern that, given the low annual incidence of infections, single-pathogen agents might not yield a sufficient return on investment to support the development costs. A recent study examined the cost effectiveness of a novel, pathogen-specific agent targeting carbapenem-resistant *A. baumannii* (CRAB) in comparison with the standard-of-care (Spellberg 2013). The analysis based on annual incidence of sensitive versus resistant infections, cost of treatments, and mortality of infections showed that a single-pathogen agent could provide benefit at a cost well below the traditional benchmarks used to define cost effective therapy. Our compounds fit the profile for a cost-effective single pathogen treatment paradigm for *Ng*

## METHODS

### Antimicrobial Susceptibility Tests for *Ng* and *Nm*

PTC-565, PTC-847, and PTC-672 susceptibility testing for *Ng* and *Nm* and other Gram-negative pathogens and normal gut organisms was performed in accordance with the CLSI M07-A9 guideline (CLSI 2012). The direct colony suspension procedure was used when testing *Ng* and *Nm*. *Ng* were tested on a gonococcal typing (GC) agar base with 1% defined growth supplement and *Nm* were tested on Mueller-Hinton Agar (MHA) supplemented with 5% sheep blood. Colonies from an overnight chocolate agar culture plate were suspended in 0.9% phosphate buffered saline (PBS), pH 7.0, to a McFarland standard. The agar plates were inoculated within 15 min after adjusting the turbidity and incubated at 37°C in an atmosphere of 5% CO_2_ for 20 to 24 h before reading the MIC values.

### Bacterial Time-dependent Kill and Post Antibiotic Effect Curves

The time-dependent kill and post antibiotic effect kinetic assays were performed in accordance with the CLSI M26-A guideline (CLSI M26-A 1999). Aliquots from the cultures at the indicated time points were serially diluted 10-fold in PBS (10^−1^ to 10^−6^) which were then spotted onto agar plates for 24 h and the colony forming units per mL (CFU/mL) determined in duplicate. The kill kinetics were represented graphically by plotting the log_10_ CFU/mL against time at each concentration of the compound.

### DNA and Protein Synthesis Assay

The DNA and protein synthesis assays were performed as described (Jyssum 1979). *Ng* 13477 was grown overnight from a single colony in 20 mL fastidious broth (FB) liquid media at 37°C with 5% CO_2_. The culture was then again diluted ~10 fold in FB and grown for 3 h to an OD of 0.4.

For protein assays, 23 mL of this culture was sedimented and washed in 50 mL minimal media without leucine [10 mM NaH_2_PO_4_ pH 7.0, 12 mM KCl, 6 mM MgCl_2_, 16 mM (NH_4_)_2_SO_4_, 24 mM NaCl, 1x BD BBL™ IsoVitaleX™ Enrichment (Thermo Fisher Scientific, Waltham MA), and 100 μM amino acids minus leucine]. The cells were then resuspended in 22 mL of this media. For the DNA synthesis assay, the cell culture was used directly without refreshing the media.

Aliquots (200 μL/well) of these cultures were transferred to 96 well microplates. To each well, 5 μL of various 40x concentrations of PTC-847 in 100% DMSO and a control with 100% DMSO were added (2.5% DMSO final). Then the [^14^C] leucine or [^14^C] uracil (1 μCi/mL), were added and the plates incubated at 35°C with 5% CO_2_ atmosphere for 3 h. At the end of the incubation, 90 μL was transferred to the wells of a 96 well filter plate containing an equal volume of 20% tricholoracetic acid (TCA). Another 90 μL was transferred to the wells of a 96 well polypropylene plate containing 10 μL of 3.5 N KOH at 37°C.

In the former case, protein and total nucleic acids were TCA precipitated at 4°C for 30 min and the resulting precipitate was collected on filters by centrifugation at 3000 x g for 6 min. In the latter case, RNA was hydrolyzed by KOH treatment at 37°C overnight and 90 μL of each hydrolysate was transferred to a 96 well filter plate containing an equal volume of 20% TCA (at 4°C). The remaining DNA was then TCA precipitated at 4°C for 30 min and collected on filters by centrifugation.

In both workups, the filters were washed with 200 μL ice cold 10% TCA and then dried. The plates were sealed, scintillation cocktail (50 μL/well) was added, and the radioactivity analyzed by scintillation counting (Perkin Elmer).

### *Ng* topoisomerase IV expression and purification

Earlier purification attempts of individual subunits lead to solubility issues of gyrase B, and we therefore chose to co-express the subunits for purification. Plasmids (pET-Duet1) containing the S-tagged *Ng parC* and His-tagged *parE* genes (cloned into sites NcoI/PstI and XhoI/BglII, respectively)were transformed into the recombination-deficient *Ec* (recA) expression strain BLR(DE3) (Novagen) to avoid gene rearrangements. The ParC and ParE proteins were induced and purified using a previously published method (Pan 1999).

Single colonies were picked from plates and grown overnight at 37°C in 50 mL of Luria-Bertani (LB) medium containing the selective antibiotic. A culture (10 mL) of the overnight growth was used to inoculate 450 mL of LB medium containing ampicillin (100 mg/mL) or kanamycin (50 mg/mL). Cells were grown at 30°C for 3 to 4 h, until the optical density at 600 nm reached 0.4 to 0.6. The culture was transferred to 18°C and IPTG was added to a final concentration of 0.5 mM, and growth was continued overnight. Bacteria were harvested by centrifugation at 5,000 x g for 15 min at 4°C. The supernatant was discarded, and the bacterial pellet was stored at −80°C.

The suspension was thawed on ice, and the pellet was resuspended in 20 mL of buffer A (20 mM Tris-HCl [pH 7.9], 500 mM NaCl, 5 mM Imidazole, containing EDTA-free complete protease inhibitor (Roche)). Lysozyme (Sigma) and Triton X-100 was added to achieve a final concentration of 0.02% and 0.1% respectively. Incubation was continued on ice for another 30 min and then the mixture was sonicated and centrifuged at 35,000 x g for 60 min.

The supernatant was carefully removed to a 50-mL sterile tube and mixed with 2 mL volume of 50% Ni-NTA resin (Qiagen) which was pre-equilibrated in the buffer A. The tube was shaken gently on a Nutator overnight at 4°C and then sedimented to remove the Ni-NTA flow through. The Ni-NTA resin was then loaded into a column and washed initially with 20 mL of buffer A, followed by washes with 20 mL of Buffer B (Buffer A plus 50 mM Imidazole). The tagged ParC and ParE protein was sequentially eluted with 20 mL buffer C (Buffer A plus 200 mM imidazole) collected as fractions that were stored frozen at −80 °C. The column fractions were examined by Western blot. The fractions containing the desired protein were pooled and dialyzed twice for 4 h against 4L of 20 mM Tris-HCl (pH 7.9), 250 mM NaCl, 1 mM DTT and 10% glycerol. The dialyzed solution was spun in a microcentrifuge at 16,000 x g for 10 min at 4°C to remove any precipitate. Purified combined topoisomerase IV was flash frozen in aliquots and stored at −80°C. The activity of *Ng* topoisomerase IV was measured using a kDNA decatenation assay and was comparable to commercially available *Ec* topoisomerase IV.

### *Ng* gyrase isolation was carried by a modification from Gross (2002)

S-tagged *gyrA* and His-tagged *gyrB* were cloned into NcoI/PstI and XhoI/BglII, sites respectively in pET-Duet1 and transformed into BLR(DE3). Growth and lysozyme treatments were as described above for topoisomerase IV (Pan 1999) and the supernatant was mixed with Ni-NTA resin and shaken overnight at 4°C. The resin was loaded into a column and the flow through containing S-tagged gyrase A was further purified using an S-protein agarose (Sigma) by elution with 20 mM Tris pH 7.5, 300 mM NaCl, 3M MgCl_2_.

His-tagged gyrase B was loaded into a Ni-NTA column, washed with 20 mM Tris-HCl [pH 8.0], 300 mM NaCl, 10% glycerol, 0.1% Triton X-100), eluted in the same buffer containing 200 mM imidazole. The purified protein judged by SDS PAGE were pooled and diluted 32-fold in buffer D (50 mM Tris HCl [pH 8.0], 2 mM dithiothreitol (DTT), 1 mM EDTA, 10% glycerol).

Further purification of gyrase A and gyrase B used a mono-Q (GE Healthcare) column in buffer D and a linear gradient elution from 0 to 1 M NaCl. Fractions were analyzed by SDS PAGE and Coomassie blue staining.

The fractions containing the proteins were pooled, concentrated and desalted using Ultrafree-15 Biomax-100 membrane centrifugal filter units (Millipore) in 10 mM Tris pH 7.9, 50 mM KCl, 0.1 mM EDTA, 2mM DTT. Glycerol was added to the purified protein at a final concentration of 25% and flash frozen in liquid nitrogen, and stored at −80°C. The protein concentrations were determined by A280 nM and calculated extinction coefficients of *Ng* GyrA (56,400 M^−1^ cm^−1^) and *Ng* GyrB (51,520 M^−1^ cm^−1^). Active gyrase (A_2_B_2_) was formed prior to the start of an assay using equal molar amounts (15 μM GyrA and GyrB).

### Bacterial DNA gyrase assays

Microplate based supercoiling assays for *Ec* or *Ng* DNA gyrase were performed as described (Maxwell 2006). *Ec* DNA gyrase was purchased (TopoGEN, Buena Vista, CO). Pierce™ black streptavidin coated 96-well microplates (Thermo Fisher Scientific, Waltham, MA) were rehydrated and washed 3x in wash buffer (20 mM Tris-HCl, pH 8.0, 137 mM NaCl, 0.01% bovine serum albumin, 0.05% Tween-20). 100 μL of 500 nM biotinylated triplex forming oligonucleotide (TFO1) was immobilized onto the streptavidin plate. Excess oligonucleotide was washed off using the wash buffer. Enzyme reactions (30 μL) containing 1 μg relaxed plasmid pNO1 DNA (Inspiralis Limited, Norwich, UK) in 35 mM Tris-HCl (pH 7.5), 24 mM KCl, 4 mM MgCl_2_, 2 mM DTT, 1.8 mM spermidine, 1 mM ATP, 6.5% glycerol, 0.1 mg/ml albumin, and 1 U of *Ec* DNA gyrase (TopoGEN, Buena Vista, CO) or DNA gyrase purified as described above from the *Ng* 13477 strain was incubated at 37°C for 30 min. 100 μL of TF buffer (50 mM sodium acetate, pH 5.5, 50 mM NaCl, 50 mM MgCl2•6 H_2_O) was then added to the reaction and the entire mixture transferred to the microplate wells after the TFO1 immobilization process. The microplate was incubated at room temperature for 30 min to allow triplex formation. Unbound plasmid was washed off with 3 x 200 μL of TF buffer, and 200 uL of 1X SYBR Gold (Invitrogen, Carlsbad, CA) in 10 mM Tris-HCl (pH 8.0), 1 mM EDTA was added and allowed to stain for 20 min. Fluorescence was read using the EnVision Multimode Plate Reader (Perkin-Elmer, Waltham, MA).

### Human DNA topoisomerase II and bacterial DNA topoisomerase IV decatenation assays

These topoisomerases can convert the large network of concatenated kinetoplast DNA (kDNA) from *C. fasciculata* DNA into decatenated DNA. Topoisomerase decatenation assays are based on the principle that decatenated DNA can be separated from concatenated DNA by microfiltration and quantified by fluorescence. Decatenation assays using purified human DNA topoisomerase II (ProFoldin, Hudson, MA), *Ec* DNA topoisomerase IV (ProFoldin, Hudson, MA), and *Ng* DNA topoisomerase IV purified in-house from the *Ng* 13477 strain as described were performed according to ProFoldin’s protocols for microplate-based topoisomerase DNA decatenation assays using kDNA (ProFoldin, Hudson, MA) and in-house reagents. The separated decatenated product was stained using the Quant-iT™ PicoGreen™ dsDNA Assay Kit (Invitrogen, Carlsbad, CA) followed by fluorescence quantitation on an EnVision Multimode Plate Reader.

### Nucleotide pools measured in *Ng*

*Ng* 13477 and PTC-847^R^ were grown overnight in FB media and then subcultured. Two *Ng* 13477 and one PTC-847^R^ subcultures were allowed to grow for 4 hours. Then PTC-847 (0.05 μg/ml) was added to one of the *Ng* 13477 subcultures and incubated an additional 1 hour at 37°C with CO_2_. The bacteria were collected by centrifugation for 15 min at 4,000 x g at 4°C. The pellets were resuspended in 5 mL cold acidified ACN (65% acetonitrile, 35% water, 100 mM formic acid) and extracted on ice for 30 min with periodic mixing. The extractions were sedimented for 20 min at 30,000 x g at 4°C. The supernatants were collected, frozen on dry ice, lyophilized to dryness, and subjected to LC/MS. ^13^C9,^15^N3-CTP was added for LC/MS analysis and peak areas of NTP and dNTPs were normalized to the peak area of ^13^C9,^15^N3─CTP (Chen 2009)

### Cloning and purification of *Ng* RNR subunits

The gene for the α subunit of *Ng* Ia RNR was optimized for expression in *Ec* and cloned into a vector with an N-terminal pET(His)_6_SUMO-Kan-N. The protein was expressed, purified to homogeneity by Ni-affinity chromatography, and the tag cleanly removed using the SUMO protease (Parker 2018). Subunit β was cloned and expressed with an N-terminal hexahistidine (His)_6_-tag that resulted in homogeneous protein with variable amounts of metallo-cofactor. The diferric tyrosyl radical cofactor was self-assembled with Fe^2+^ and O_2_ to give 0.7 Y•s/β_2_. *Ng* His6-tagged RNR subunits as previously described (Ravichandran 2020, Greene 2020).

### Activity assay of *Ng* RNR

To optimize RNR activity a 1:1 ratio of α_2_ and β_2_ subunits was examined over a physiological concentration range of 0.01 to 10 μM. Its specific activity (SA) determined in a reaction mix of 100 μM *Ec* TR, 1 μM *Ec* TRR, and 0.2 mM NADPH at 37 °C. The SA for 0.01 to 0.12 μM subunits was measured with 1 mM GDP, 0.25 mM TTP, and 0.2 mM NADPH using the continuous spectrophotometric assay. The SA for 0.2 to 10 μM was measured with 1 mM 5-[^3^H]-CDP, 3 mM ATP and 2 mM NADPH by the discontinuous radioactive assay. Assay detection methods followed the standard protocols (Ge 2003, Ravichandran 2020).

The assay for inhibition of *Ng* RNR by PTC-847 and PTC-672 was performed spectrophotometrically using the GDP/TTP protocol with the addition of various concentrations of PTC-847 (0.1 to 8 μM) or PTC-647 (0.025 to 2.5 μM). The inhibition of RNR by dATP was performed spectrophotometrically with the addition of various concentrations of dATP (2 nM to 1 μM). At concentrations of dATP from 50 to 200 μM, [3H]-CDP/ATP and the discontinuous assay was used.

### Protocol for Negative Stain EM

Negative stain EM samples contained 1:1.5 α:β, 15 ng/μL protein. Concentrations of α and β were determined using A280 nm (ε = 89050 M^−1^cm^−1^ for α and 61310 M^−1^cm^−1^ for β). αs (WT and mutants) and β were first combined on ice at 150 ng/μL and incubated for one min in the presence of specified nucleotides, also at 10x concentration. The samples were then diluted tenfold in RNR assay buffer (50 mM HEPES pH 7.6, 15 mM MgCl_2_, 1 mM EDTA) and 5 μL of the mixture was applied to carbon-coated 300 mesh Cu grids (Electron Microscopy Sciences) after glow discharging for 1 min at −15 mA in an EasiGlo glow discharger (Ted Pella). The protein was adsorbed to the grid for 1 min and excess liquid was removed by blotting. The grid was stained 3x with 5 μL of 2% uranyl acetate (VWR) with blotting immediately after application for the first two rounds of staining. The final uranyl acetate stain was allowed to sit on the grid for 45 s before blotting. All blotting was completed manually with filter paper (Whatman, grade 40).

Screening images for the *Ng* RNR negative stain reconstruction were taken on a Tecnai Spirit (FEI) instrument with a XR16 camera (AMT) operated at 120 kV at 68,000 × magnification. Screening images for the H25R and S41L α variant samples were taken on a Tecnai F30 (FEI) operated at 200 kV equipped with an FEI Falcon II direct electron detector. 5-10 grid squares were examined, and a representative field is shown for each protein sample. For the EM reconstruction of *Ng* RNR, images were taken on a Tecnai F20 Twin (FEI) operated at 120 kV equipped with a US4000 CCD detector (Gatan) at 1.79 Å/pix.

All processing for negative stain data sets was completed in Relion 3.0.8 (Scheres 2012). Micrographs were imported and CTF-corrected prior to manual picking of 1000 particles subdivided into 5 classes to train Relion’s autopicking algorithm. Autopicking yielded ~27000 particles for the inactivated complex. These particles were 2D classified (100 classes, 25 iterations) and an initial 3D model was created (no symmetry imposed) prior to 3D classification (1 class, 100 iterations, D2 symmetry) and 3D auto-refinement. Auto re-finement was completed on 12469 particles and yielded a structure at 21 Å resolution (0.143 FSC). Following autorefinement, the inactive *Ec* RNR structure (PDB 5CNS) was docked into the *Ng* RNR EM map using the Chimera MaptoModel functionality (Pettersen 2004, Zimanyi 2016).

### In vivo studies using a mouse model of vaginal *Ng* infection

All animal studies were performed under IACUC approved protocols at AAALAC-certified animal facilities. Female ovariectomized Balb/c mice (Charles River, Wilmington, MA) were obtained at 3 weeks of age and acclimatized for ~1 week prior to initiation of a fitness or efficacy study. Day 0 was defined as the day mice were inoculated vaginally with bacteria suspended in saline. A single 17-β-estradiol pellet (0.5 mg, 21-day release) was then implanted in mice with a trochar on Day −2. Mice were then treated with three antibiotics to control commensal flora induced by the estradiol. Vancomycin HCl (0.6 mg) and streptomycin sulfate (0.3 mg) were administered intraperitonealy and trimethoprim sulfate (0.8 mg) was administered by oral gavage twice daily from Day −2 to Day 1 of study. On Day 2 of the study the administration of all antibiotics except streptomycin sulfate was stopped. Streptomycin sulfate was then administered daily from Day 2 throughout the remainder of the study.

### Fitness testing of PTC-672 resistant *Ng* in the mouse model of vaginal *Ng* infection

The relative fitness cost associated with PTC-672 resistance was determined in vivo by competition experiments using the validated mouse model of vaginal *Ng* infection described above. This study evaluated the relative fitness of the PTC-672 resistant strains Iso 13477 Strep^R^-PTC-672^H25R^ and Iso 13479 Strep^R^-PTC-672^H25R^ in competition with the isogenic PTC-672 susceptible parent strains Iso 13477 Strep^R^ and Iso 13479 Strep^R^. On Day 0 of the study, mice pre-treated as described above were inoculated vaginally with either the Strep^R^ parent strain, the isogenic Strep^R^-PTC672^H25R^ strain, or a mixture of the Strep^R^ parent and the isogenic Strep^R^-PTC672^H25R^ strains suspended in saline and swabbed daily over a one week period. At each sampling time point, the numbers of total (susceptible + resistant bacteria) and resistant bacteria were determined by plating onto non-selective and PTC-672-supplemented plates, respectively. Relative fitness was expressed as the CI, calculated as the ratio of bacterial burdens between the PTC-672 resistant and susceptible strains at each time point divided by the baseline ratio at the beginning of the experiment.

### In vivo efficacy studies using the mouse model of vaginal *Ng* infection

The efficacy of an increasing single oral dose of PTC-672 administered by oral gavage was evaluated in the mouse model described above. The antibiotics, ciprofloxacin and ceftriaxone, were used as positive controls for efficacy. Efficacy was defined as sustained clearance of infection for 5 days post dosing in 10/10 mice in a treatment group.

On Day 0 of the study, mice pre-treated as described above were inoculated vaginally with Iso 13477 Strep^R^ or Iso 13479 Strep^R^ bacteria suspended in saline. On Day 2 mice were randomized into treatment groups based on Day 1 CFU counts. All mice were treated on Day 2 with a single oral dose of either vehicle comprising 0.5% hydroxypropyl methylcellulose and 0.1% Tween^®^ 80, ciprofloxacin, ceftriaxone, or PTC-672. In the Iso 13477 Strep^R^ portion of the study, PTC-672 was tested at doses of 10, 15, 20, 25, and 30 mg/kg and ciprofloxacin at 30 mg/kg. In the Iso 13479 Strep^R^ portion of the study, PTC-672 was tested at doses of 30 and 60 mg/kg and ciprofloxacin at 100 mg/kg. Mice were vaginally swabbed daily using a sterile swab over a one-week period. At each sampling time point, the numbers of resistant bacteria were determined by plating onto PTC-672-supplemented plates and the level of infection was determined for each test mouse in each treatment group.

## ACKNOWLEGEMENTS

This project had been funded in part by the Wellcome Trust through a Seeding Drug Discovery award (097753) and by the National Institute of Health Grants R35 GM126982 (CLD) and NIH Pre-Doctoral Training grant T32 GM007287 (TSL) and NSF GRFP 2017246757 (TSL). C.L.D. is a Howard Hughes Medical Institute Investigator. Chang Cui was supported by NIH grant GM47274 to Daniel G Nocera, Department of Chemistry at Harvard University and NIH grant GM29595 (JS). The authors wish to thank Richard Sheridan for assembling the data and getting the manuscript in submittable format.

