## Supplementary Data for "Ribonucleotide reductase, a novel target for gonorrhea"

**Current Affiliations:**

Melissa Dumble: PMV Pharmaceuticals, 8 Clarke Drive, Cranbury, NJ, 08512 USA.

^§^Aleksey Gerasyuto: Schrödinger, Inc., 120 West 45th Street, 17th Floor, New York, NY 10036

^§^Jiashi Wang: Schrödinger, Inc., 120 West 45th Street, 17th Floor, New York, NY 10036

^Ψ^Suzanne Letinski: Bristol Myers Squibb, 1 Squibb drive, New Brunswick, NJ 08901

**SUPPLEMENTARY INFORMATION**

| **Antibiotic / Compound** | **MIC range**  **(µg/mL)** | | **Value for decreased susceptibility (µg/mL)** | **Value for elevated MICs (µg/mL)** | **Isolates with elevated MICs** |
| --- | --- | --- | --- | --- | --- |
| Azithromycin | ≤0.03 to >256 | >8500-fold | ≥2 | Na | 9 |
| Cefixime | ≤0.002 to 4 | >2000-fold | ≥0.5 | ≥0.25 | 16 |
| Ceftriaxone | ≤0.002 to 1 | >500-fold | ≥0.5 | ≥0.12 | 5 |
| Ciprofloxacin | ≤0.03 to >16 | >500-fold | ≥0.12 | Na | 71 |
| Tetracycline | <0.5 to >16 | >32-fold | ≥2 | Na | 179 |
| PTC-847 | 0.03 to >1 | >32-fold | - | >1 | 2 |
| PTC-672 | 0.015 to 0.25 | 16-fold | - | - | 2 |

**Table S1. PTC-847 and PTC-672 MIC_90_ values across 206 *N. gonorrhoeae* strains collected at the Public Health England.** Two isolates gave elevated PTC-847 or PTC-672 MIC values. However, these isolates were sensitive to all other antibiotics. Due to the selectivity of the PTC compounds, these two strains will be genotyped to confirm they are *Neisseria* species. PTC-compound susceptibility testing was performed in accordance with the Clinical and Laboratory Standards Institute (CLSI) M07-A9 guideline (CLSI M07-A9 2012).

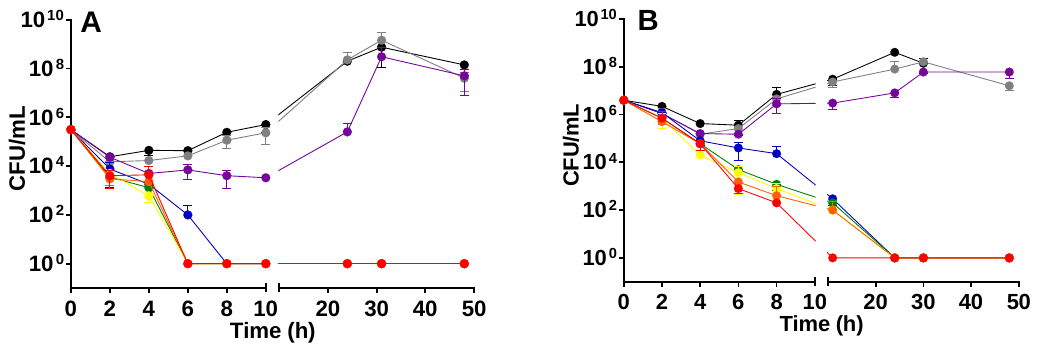

**Figure S1. Time -dependent kill of *Ng* upon PTC-847 (A) or PTC-672 treatment (B).** In both graphs A and B, the DMSO control data is shown in black, data for 0.25 x MIC, 0.5 x MIC 1 x MIC, 2 x MIC, 4 x MIC, 8 MIC and 16 x MIC are shown in grey, purple, blue, green, yellow, orange and red respectively. At PTC-847 or PTC-672 concentrations of ≥ 1 x MIC, a >3-log reduction in measured *Ng* 13477 colony forming units per milliliter of culture (CFU/mL) was observed. The time-dependent kill assays were performed in accordance with the CLSI M26-A guideline (CLSI M26-A 1999).

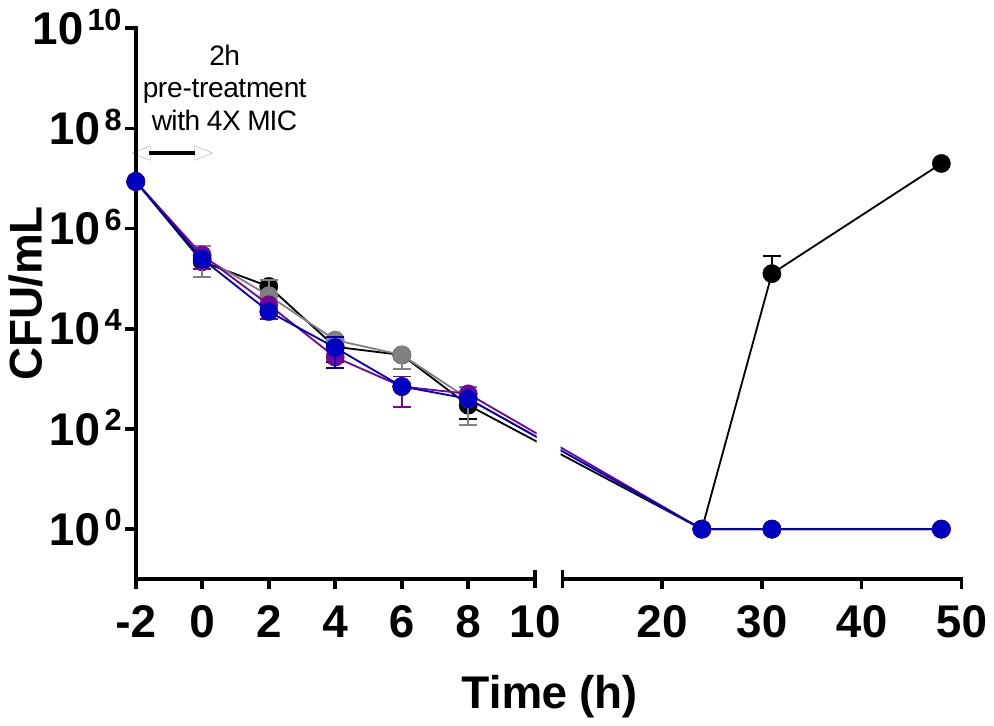

**Figure S2. Time-dependent post antibiotic effect observed with PTC-847.** *Ng* 13477 was pre-treated with PTC-847 at 4X MIC for 2h then allowed to grow in culture media supplemented with PTC-847 at 0, 0.25, 0.5, or 1X MIC (Data for 0, 0.25 x MIC, 0.5 x MIC, and 1x MIC are indicated in black, grey, purple, and blue respectively). A >6-log reduction in measured *Ng* 13477 CFU/mL was observed at 20 h post pre-treatment in the absence of PTC-847. The post antibiotic effect assay was performed in accordance with the CLSI M26-A guideline (CLSI M26-A 1999).

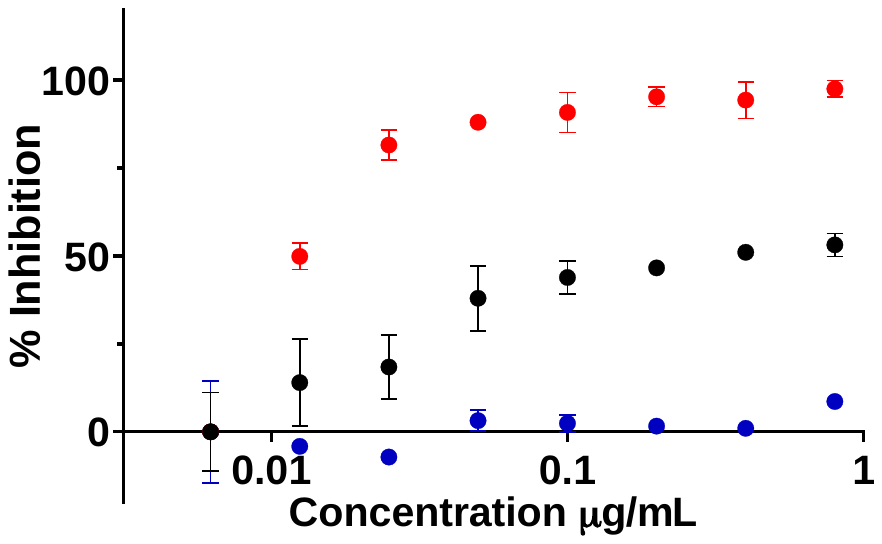

**Figure S3. PTC-847 inhibition of DNA synthesis in *N. gonorrhoeae* strain 13477.** We followed the incorporation of the radiolabeled precursors into total nucleic acids (black circles), DNA (red circles), and protein (blue circles). In *Ng*, only radiolabeled uracil is incorporated into DNA or RNA, which was also shown to be the case for *Nm* (Jyssum 1979).

| Strain | MIC fold | Resistance Frequency | | |
| --- | --- | --- | --- | --- |
|  |  | PTC-847 | Ciprofloxacin | Ceftriaxone |
| *Ng* 13477 | 32X | 5.6 x 10^-9^ | ≤8 x 10^-9^ | ─ |
|  | 16X | 5.6 x 10^-9^ | ≤8 x 10^-9^ | ─ |
|  | 8X | 1.1 x 10^-8^ | ≤8 x 10^-9^ | ≤1.9 x 10^-9^ |
|  | 4X | 3.6 x 10^-8^ | 1.2 x 10^-8^ | 7.7 x 10^-9^ |

**Table S2. Frequency of resistance to PTC-847.** By plating at high cell density on agar plates containing PTC-847 at 4, 8, 16 and 32-fold MIC, Ng 13477 exhibited a spontaneous or acquired frequency of resistance to PTC-847 on the order of 10^-8^. The colonies obtained were passaged multiple times on PTC-847-containing plates to obtain a stable PTC-847-resistant strain (PTC-847^R^).

| Inhibitor  type | Class | Compound / Antibiotic | *Ng* MIC (µg/mL) | |
| --- | --- | --- | --- | --- |
|  |  |  | Wild-type | PTC-847^R^ |
| DNA | novel | PTC-847 | 0.05 | 15.6 |
|  | fluoroquinolones | ciprofloxacin | 0.003 | 0.003 |
|  |  | moxifloxacin | 0.003 | 0.003 |
|  |  | enrofloxacin | 0.004 | 0.004 |
|  |  | delafloxacin | 0.05 | 0.05 |
| Protein | oxazolidinone | linezolid | 2 | 2 |
|  | aminoglycosides | gentamicin | 2 | 2 |
|  |  | kanamycin | 6.2 | 6.2 |
|  | macrolides | solithromycin | 0.1 | 0.05 |
|  |  | erythromycin | 0.2 | 0.2 |
|  | tetracyclines | tetracycline | 1 | 1 |
|  |  | tigecycline | 0.4 | 0.4 |
| RNA | rifamycin | rifampicin | 0.1 | 0.1 |
| Cell wall | beta-lactam | ampicillin | 0.06 | 0.06 |
|  | cephalosporins | cefepime | 0.002 | 0.002 |
|  |  | ceftriaxone | 0.0002 | 0.0004 |

**Table S1. PTC-847^R^ strain exhibits no cross resistance to existing antibiotics.** Susceptibility of the WT *Ng* 13477 strain and the PTC-847^R^ strain were measured for a wide variety of antibiotics having different modes of action. MICs were determined in accordance with the CLSI M07-A9 guideline (CLSI M07-A9 2012). The PTC-847^R^ strain was equally sensitive to all classes of antibiotics as the susceptible WT *Ng* 13477 strain, except for resistance to PTC-847. The PTC-847 MIC for WT *Ng* 13477 was 0.05 µg/mL compared to 15.6 µg/mL for the PTC-847^R^ strain.

| NTP/dNTP Ratio | Treatment | | Fold change |
| --- | --- | --- | --- |
|  | DMSO | PTC-847  (1X MIC) |  |
| CTP/dCTP | 10.91 | 33.79 | 3.1 |
| UTP/dTTP | 7.63 | 24.12 | 3.2 |
| ATP/dATP | 17.4 | 44.2 | 2.5 |
| GTP/dGTP | 19.9 | 24.9 | 1.3 |

**Table S4. Nucleotide pools measured in WT *Ng* 13477.** The susceptible WT Ng 13477 strain was grown to log phase and treated with DMSO or PTC-847 at 1X MIC for 1 h. Nucleotides were extracted in acidified acetonitrile/H_2_O (65:35) followed by centrifugation. The supernatant was lyophilized and subjected to LC/MS. ^13^C9,^15^N3─CTP was added for LC/MS analysis. Peak areas of NTP and dNTPs were normalized to the peak area of ^13^C9,^15^N3─CTP.

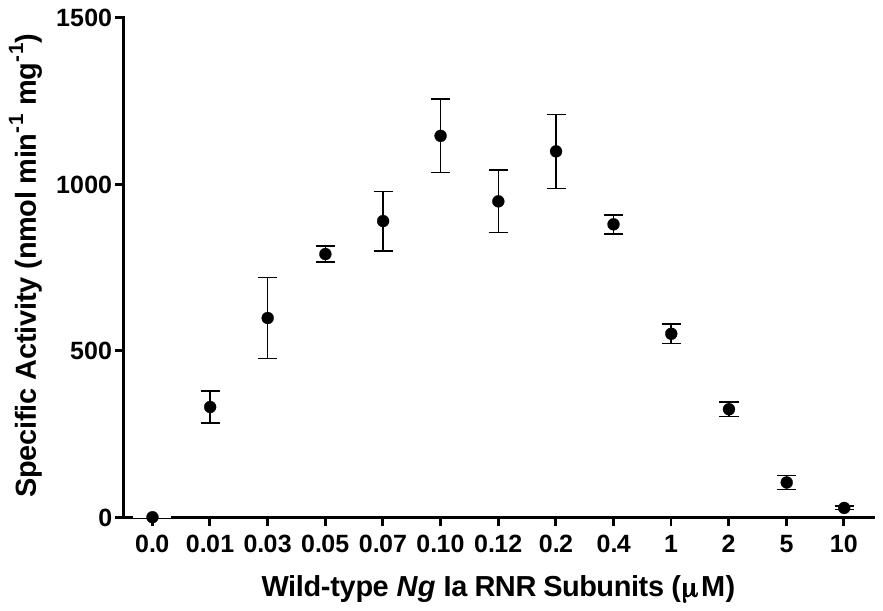

**Figure S4. Activity assay of *Ng* RNR.** To optimize RNR activity, a 1:1 ratio of α_2_ and β_2_ subunits was examined over a physiological concentration range of 0.01 to 10 µM and its specific activity (SA) determined in a reaction mix of 100 µM *Ec* thioredoxin (TR), 1 µM *Ec* thioredoxin reductase (TRR), and 0.2 mM NADPH at 37 °C. The SA for 0.01 to 0.12 µM subunits was measured with 1 mM GDP, 0.25 mM TTP, and 0.2 mM NADPH using a spectrophotometric assay whereas the 0.2 to 10 µM SA was measured with 1 mM 5-[^3^H]-CDP, 3 mM ATP and 2 mM NADPH by radioactive assay. Fitting the data from 0.01 to 0.12 µM using $v=\frac{V_{max}[Subunits]}{\left[ Subunits \right]+K_{m}}$ gave *V*_max_ = 1300 nmol min^-1^ mg^-1^ and *K*_m_ = (0.030±0.006) µM.

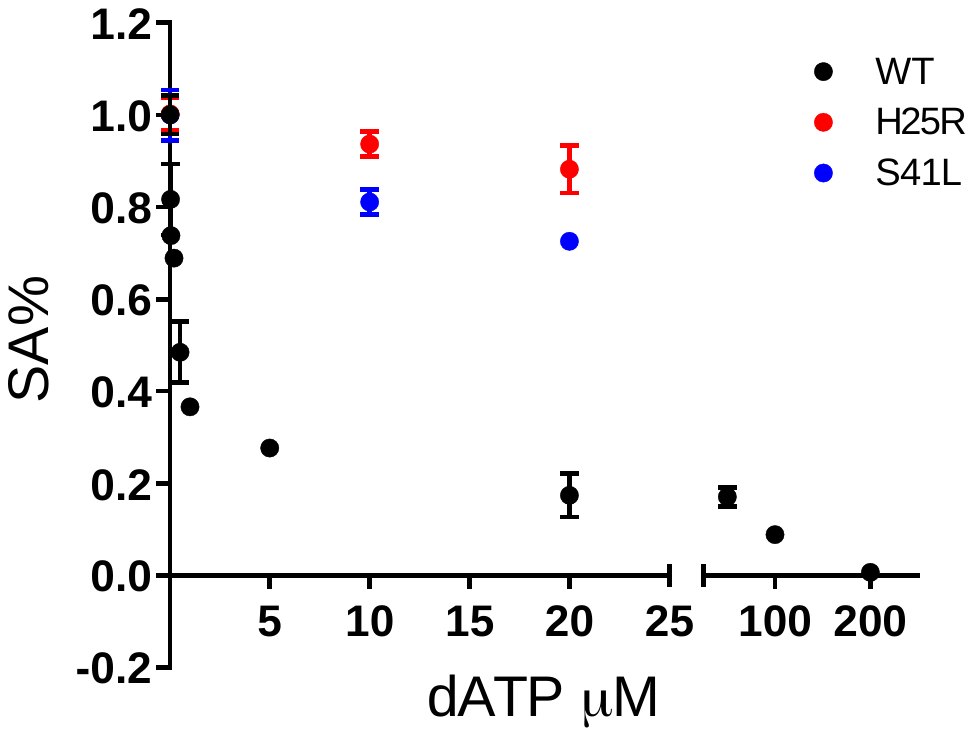

**Figure S5. dATP inhibition of *Ng* RNR.** The continuous spectrophotometric assay with GDP/TTP (Figure S4) was used for concentrations of dATP to 25 μM and NADPH concentration was 0.2 mM. The discontinuous radioactive assay with [5-^3^H] CDP (1 mM) and ATP (3 mM) was used for dATP at 50 to 200 μM and NADPH concentration at 2 mM (see ref 38). The assay with mutant αs (H25R and S41L) used the spectrophotometric assay with GDP/TTP.

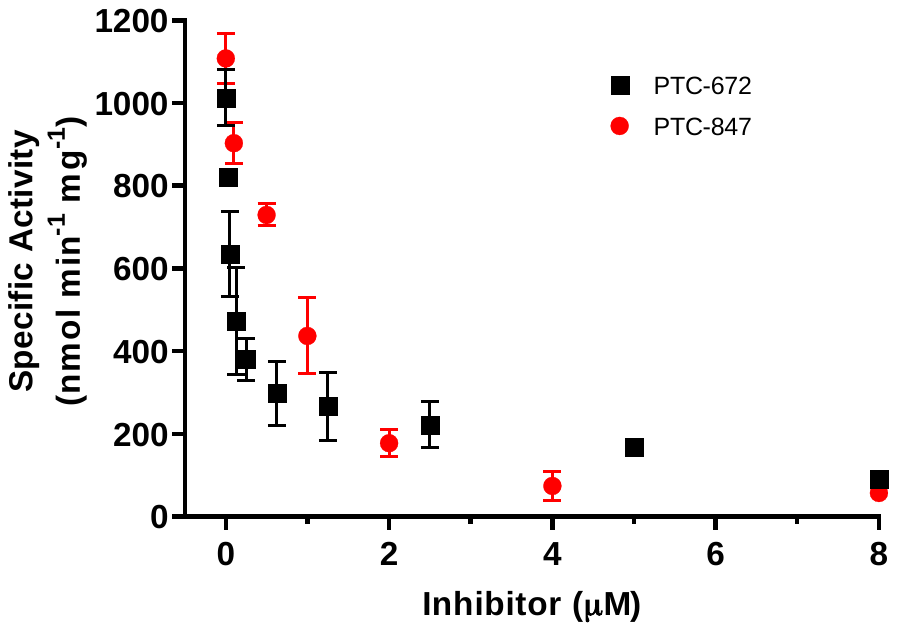

**Figure S6. Inhibition of *Ng* Ia RNR by PTC-847 and PTC-672.** A 1:1 mixture of α and β subunits was incubated with each compound and assayed for activity spectrophotometrically at 37 °C with PTC-847 (red) and PTC-672 (in black). The assay used 0.1 µM α_2_, 0.1 µM β_2_, 1 mM GDP, 0.25 mM TTP, 100 µM *Ec* TR, 1 µM *Ec* TRR, 0.2 mM NADPH and various concentrations of PTC-847 (0.1 to 8 µM) or PTC-672 (0.025 to 8.0 µM).

| Compound | MIC (µg/mL) | | | |
| --- | --- | --- | --- | --- |
|  | *Bacteroides fragilis* | *Bifidobacterium bifidum* | *Bifidobacterium longum* | *Clostridioides difficile* |
| Solithromycin | 1 | <0.06 | <0.06 | <0.06 |
| PTC-847 | 32 | 32 | 16 | 32 |
| PTC-672 | >32 | >32 | >32 | >32 |

**Table S5. PTC-672 and PTC-847 do not inhibit a representative panel of normally occurring intestinal organisms.** The panel was constructed to test for inhibition of normally occurring intestinal organisms (Thursby 2017). Susceptibility testing was performed in accordance with the Clinical and Laboratory Standards Institute (CLSI) M07-A9 guideline (CLSI M07-A9 2012).

| Compound | MIC (µg/mL) | | |
| --- | --- | --- | --- |
|  | *N. mucosa* ATCC25996^†^ | *N. polysaccharea* 43768^‡^ | *Ng* 13477 |
| Broad spectrum | 0.78 | 0.39 | 0.098 |
| AZD0914 | 0.78 | 0.39 | 0.049 |
| Ciprofloxacin | 0.008 | 0.008 | 0.004 |
| PTC-672 | 1.56 | 0.39 | 0.05 |

**Table S6. Commensal *Neisseria* are less sensitive to PTC-672 than *Ng* 13477.** Two commensal *Neisseria* ‘type strains’ swabbed from the oropharynx of healthy volunteers (^†^Berger 1971 and ‡Riou 1987) were tested for susceptibility to inhibition in accordance with the Clinical and Laboratory Standards Institute (CLSI) M07-A9 guideline (CLSI M07-A9 2012).
